# HuMMANet: A Harmonized Cross-Study Resource for Integrative Analysis of Human Gut Microbiome–Metabolome Associations

**DOI:** 10.64898/2026.08.24.746727

**Authors:** Shivangi Verma, Nalin Arora, Chandra Prakash Ajay, Pankhuri Singh, Himel Mallick, Tarini Shankar Ghosh

## Abstract

Deciphering gut-microbiome–to–host-metabolome interaction is critical for understanding how microbial communities generate bioactive signals that shape host physiology and disease. Progress, however, has been hindered by inconsistent metabolite annotations, poor interoperability across studies, and the absence of integrated resources placing microbiome-derived metabolites within their functional, microbial, physiological, and clinical context. Here we present HuMMANet (Human Microbiome–Metabolome Annotation Network), a harmonized resource integrating 46 paired gut microbiome–metabolome studies (59 study-units; 14,405 samples; 13 disease categories plus a healthy/control reference category) with a scalable metabolite-harmonization framework. HuMMANet resolves heterogeneous annotations through a multi-stage workflow spanning RefMet, HMDB, PubChem, Metabolomics Workbench, SMPDB, MiMeDB-2.0, GNPS/microbeMASST, DrugBank, and DrugCentral, yielding a reference atlas of 54,914 unique metabolites, annotated with standardized chemical identifiers, biochemical pathways, microbial producer associations, physiological distributions, disease links, and structural relationships to approved therapeutics — a unified reference framework for microbiome– metabolome research.

Applying HuMMANet to a multi-cohort integration of adult serum and fecal metabolomes, we identified 519 serum and 322 fecal metabolites reproducibly associated with gut microbial community composition (PERMANOVA, P < 0.05 in at least 50% of studies in which detected), enriched for specific biomolecular classes and pathways. Cross-referencing these against Health-Associated-Core-Keystone (HACK) taxa revealed 58 serum and 25 fecal metabolites (HACK-positive) whose taxon-level associations tracked positively with the taxon-specific-HACK indices. These reproducible metabolomic signatures of microbiome health included indole-3-propionic acid, a gut barrier-protective microbial tryptophan metabolite, and 3-phenylpropionate. Drug-similarity annotation within HuMMANet linked 16 of this serum and 13 fecal HACK-positive metabolites to therapeutics used in neurological, inflammatory, and vascular disease. Conversely, 38 serum and 65 fecal metabolites, including imidazole propionate and long-chain acylcarnitines such as ACar 18:0, showed HACK-negative signatures previously associated with dysbiosis-linked disease. GNPS/microbeMASST and MiMeDB-2.0 annotations further traced subsets of these metabolites to putative bacterial producers. HuMMANet thus provides a standardized framework for reproducible microbiome–metabolome integration, enabling cross-study discovery and translational prioritization of conserved microbiome-derived metabolic signatures across human populations and disease states.

## Introduction

The gut microbiome, the largest microbial ecosystem in the human body, encodes a functional repertoire nearly two orders of magnitude larger than the human genome^1^. Much of this capacity is devoted to metabolites that mediate microbiome-host interactions: signalling molecules and biochemical functions the human genome has not evolved to perform independently^2^, including short-chain fatty acids from fermentation of microbe-accessible carbohydrates^3^, essential vitamins^4^, neurotransmitters^5^ and metabolic compounds modulating intestinal barrier and blood-brain-barrier functions^2^. Through these molecules, gut microbes influence distant organs, with individual metabolites having been mechanistically implicated in inflammatory bowel disease, cardiometabolic disorders, colorectal cancer, neurological disease, and unhealthy ageing^2,6–9^. Deciphering these microbial metabolic pathways is therefore central to understanding microbiome-health relationships. A subset even shows structural similarity to clinically used drugs^10–12^, raising translational potential but realizing this first requires knowing which microbial metabolites are reproducibly linked to host health, and that remains unresolved.

Many studies have integrated metabolomics with microbiome profiling at the population level, in disease contexts, and in therapeutic interventions^13–15^, while also focusing on the development of machine-learning and deep-learning frameworks to identify diagnostic taxa, metabolites, and microbiome–metabolome modules linked to disease phenotypes^16–21^. Despite this progress, published paired microbiome–metabolome data remain severely underutilized. Measurements now exist for well over ten thousand samples across geographies and disease contexts, yet large-scale integration remains elusive, chiefly for computational reasons rooted in inconsistent metabolite annotation. The same molecule is reported as a free-text name in one study, a database identifier in another, and an unresolved mass-to-charge feature in a third, with no automated matching recognizing these as identical. No existing resource, to our knowledge, resolves this variability while integrating curated microbe–metabolite knowledge, so each integration effort reconciles annotations independently, without reusable output, and even reconciled metabolites still lack the pathway, microbial, physiological, and pharmacological context needed for biological interpretation. Consequently, meta-analysis remains confined to a small, arbitrary fraction of the measured metabolome, and reproducible microbiome–metabolome signatures across populations remain difficult to establish.

Existing resources address parts of these needs but remain limited in scale and fragmented across repositories. The Curated Gut Microbiome–Metabolome Dataset Collection was purpose-built for paired observations but is limited to 14 studies and ∼2,900 samples^22^. The Human Microbiome Bioactives Resource (HMBR) catalogues 15 studies and 8,790 samples^23^, though its microbiome and metabolome measurements are not linked at the sample level. The Microbial Metabolome Database (MiMeDB-2.0)^24^ has recently integrated compound entries from the Collaborative Microbial Metabolite Center knowledgebase (CMMC-KB)^25^, a community-curated resource that incorporates experimentally supported microbial metabolite annotations, including evidence from germ-free versus colonized animal models. MiMeDB-2.0 and microbeMASST^26^ similarly provide extensive curated and experimentally derived microbe–metabolite associations that remain unreconciled against each other and against the conventions used in primary studies. Recent structure-centric search infrastructure such as StructureMASST^27^ similarly enables large-scale structure-to-sample mapping across public repositories, and microbiomeMASST^28^ links microbial MS/MS spectra to harmonized study metadata across 467 public datasets and 144,424 mass spectrometry files; however, none of these resources resolve the metabolite-name inconsistencies across curated, paired human microbiome–metabolome studies that limit reproducible cross-cohort meta-analysis. Existing annotation tools share three limitations: single-dataset scope, names resolved without the chemical descriptors and cross-referenced identifiers downstream interpretation requires, and dependence on pipeline-specific inputs. MetaboAnalyst^29^, IDmapping^30^, and MetaboliteAnnotator^31^ annotate single datasets; metID^32^ is LC–MS-specific^33^; MAPS^34^ requires pre-generated outputs from MZmine^35^, GNPS2^36^, SIRIUS^37^, and MS2Query^38^; and metLinkR^39^, the only multi-study tool, standardizes names without ontology or cross-reference integration.

To address these gaps, we developed HuMMANet (Human Microbiome–Metabolome Annotation Network), a unified resource integrating paired gut microbiome–metabolome datasets. HuMMANet combines (1) a multi-stage framework that reconciles heterogeneous study-specific metabolite annotations against RefMet^40^, PubChem^41^, HMDB^42^, and the Metabolomics Workbench^43^, and (2) an integrated knowledgebase linking harmonized metabolites to biochemical pathways^44^, putative microbial producers^24,26^, physiological distribution across human biofluids^42^, disease associations^42^, and structural similarity to approved therapeutics^45,46^. Notably, we incorporate the 2026 release of MiMeDB (MiMeDB-2.0)^24^, which (as described above) integrates compound entries from the CMMC-KB^25^, ensuring HuMMANet reflects the most current community-curated microbial metabolite annotations available from the outset, rather than requiring separate downstream reconciliation. Applied to 46 studies comprising 14,405 paired samples across 13 disease categories plus a healthy/control reference category, the framework reconciled 33,469 study-specific annotations into 14,550 harmonized metabolites, which together with MiMeDB-2.0 and 60,781 LC–MS/MS files from microbial monoculture experiments yield a reference of 54,914 annotated metabolites in the final HuMMANet reference atlas^24,26,47^. Harmonization increased shared-metabolite detection by a mean of 19-fold across study-pairs and recovered shared metabolites for 583 study-pairs that previously had none. Applied to independent serum and fecal cohorts, HuMMANet identified metabolite panels reproducibly associated with gut microbial composition and graded these against Health-Associated Core-Keystone (HACK) taxa^48^, identifying indole-3-propionic acid and 3-phenylpropionate among the metabolites most consistently linked to microbiome health, both with documented physiological benefit^49,50^. Structural similarity analysis further identified related approved therapeutics. By transforming metabolite annotation from a per-study bottleneck into a reusable, expandable resource, HuMMANet provides a foundation for reproducible microbiome–metabolome discovery at population scale.

## Results

### Creation of the HuMMANet resource

To establish a foundation for cross-study microbiome–metabolome integration, we systematically collated studies reporting paired gut microbiome and host metabolome measurements from the same individuals at the same time point, identified via PubMed MeSH-based searches and manual curation (**Methods; Figure 1A**), yielding 46 studies comprising 14,405 paired observations with publicly available sample-level metadata. For methodological consistency, these were stratified into 59 study-units by analytical modality, microbiome sequencing strategy, and biological matrix (targeted/untargeted metabolomics; 16S/WGS; serum/fecal), together constituting the HuMMANet resource.

**Figure 1:**
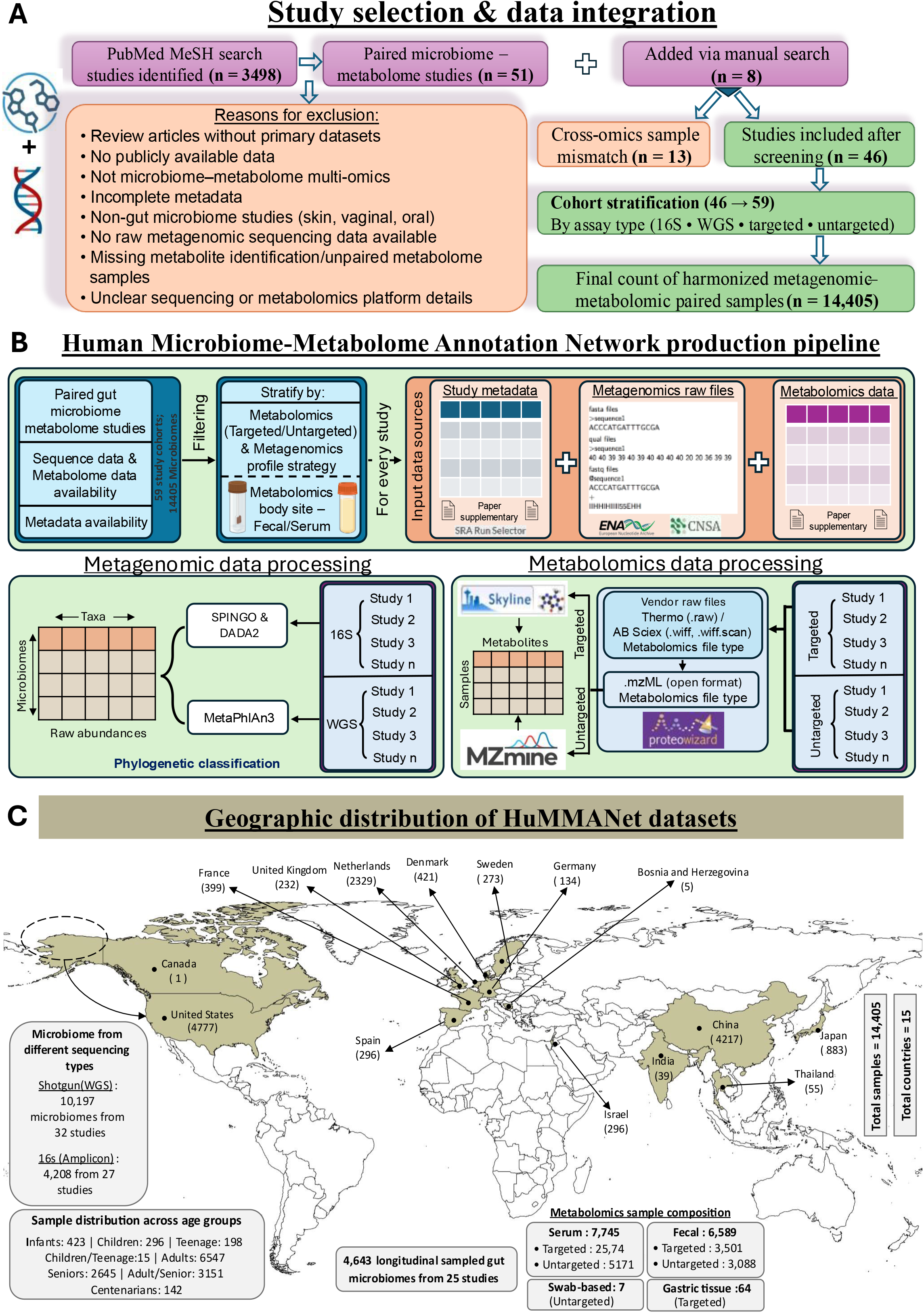
Creation of the HuMMANet resource and the integrative analysis framework. **(A)** Systematic identification, screening, and inclusion of publicly available human microbiome– metabolome studies. PubMed MeSH-based searches combined with manual curation identified candidate studies, which were filtered for public data availability, paired microbiome–metabolome measurements, and quantifiable metabolomics data (see Methods for full inclusion/exclusion criteria). After exclusion of cross-omics sample mismatches and cohort stratification by assay type, the final resource comprised 59 harmonized study-units encompassing 14,405 paired microbiome– metabolome samples from 46 studies. **(B)** Resource creation pipeline of HuMMANet. Public repositories were queried to retrieve paired microbiome and metabolomics datasets (as noted in **A**). Metabolomics data were systematically pre-processed using standardized workflows for targeted and untargeted studies, including raw data conversion, peak detection, alignment, and feature quantification, while microbiome data were processed using unified pipelines for 16S amplicon sequencing (16S) datasets using SPINGO/DADA2 and whole-metagenome shotgun sequencing (WGS) datasets using MetaPhlAn3. Processed abundance matrices, metadata, and accession-linked files were curated and harmonized across studies to generate study-wise, analysis-ready microbiome and metabolome profiles. **(C)** Global distribution of HuMMANet samples by geographic origin (bar/map; n = 14,405 samples across 15 countries), comprising WGS and 16S microbiome data paired with targeted and untargeted metabolomics. Additional panels summarize sample composition by biospecimen type, sequencing strategy, and age group.

Having assembled this collection, we next ensured cross-study comparability. All datasets underwent standardized processing and curation (**Figure 1B**). Study- and sample-level metadata were harmonized using the NCBI SRA Run Selector and study-specific supplementary information (Available at: https://www.ncbi.nlm.nih.gov/Traces/study/). Metabolite annotations were curated to retain only biologically interpretable compounds with stable chemical identities **(Table S2)**. Raw metabolomics and microbiome datasets were then processed using standardized pipelines (MZmine3, DADA2, SPINGO, and MetaPhlAn3) according to analytical modality^35,51–53^.

In its assembled form, HuMMANet includes datasets from North America, Europe, and Asia, with broad representation from the United States, China, and several European countries **(Figure 1C)**. HuMMANet represents a >4-fold expansion over the curated collection by Muller et al. **(Figure 2A)**, integrating whole-genome shotgun (10,197 samples) and 16S rRNA (4,208 samples) sequencing, broader geographic representation, 13 disease categories plus a healthy/control reference category (versus 7 previously), and expanded targeted/untargeted metabolomics coverage^22^.

**Figure 2:**
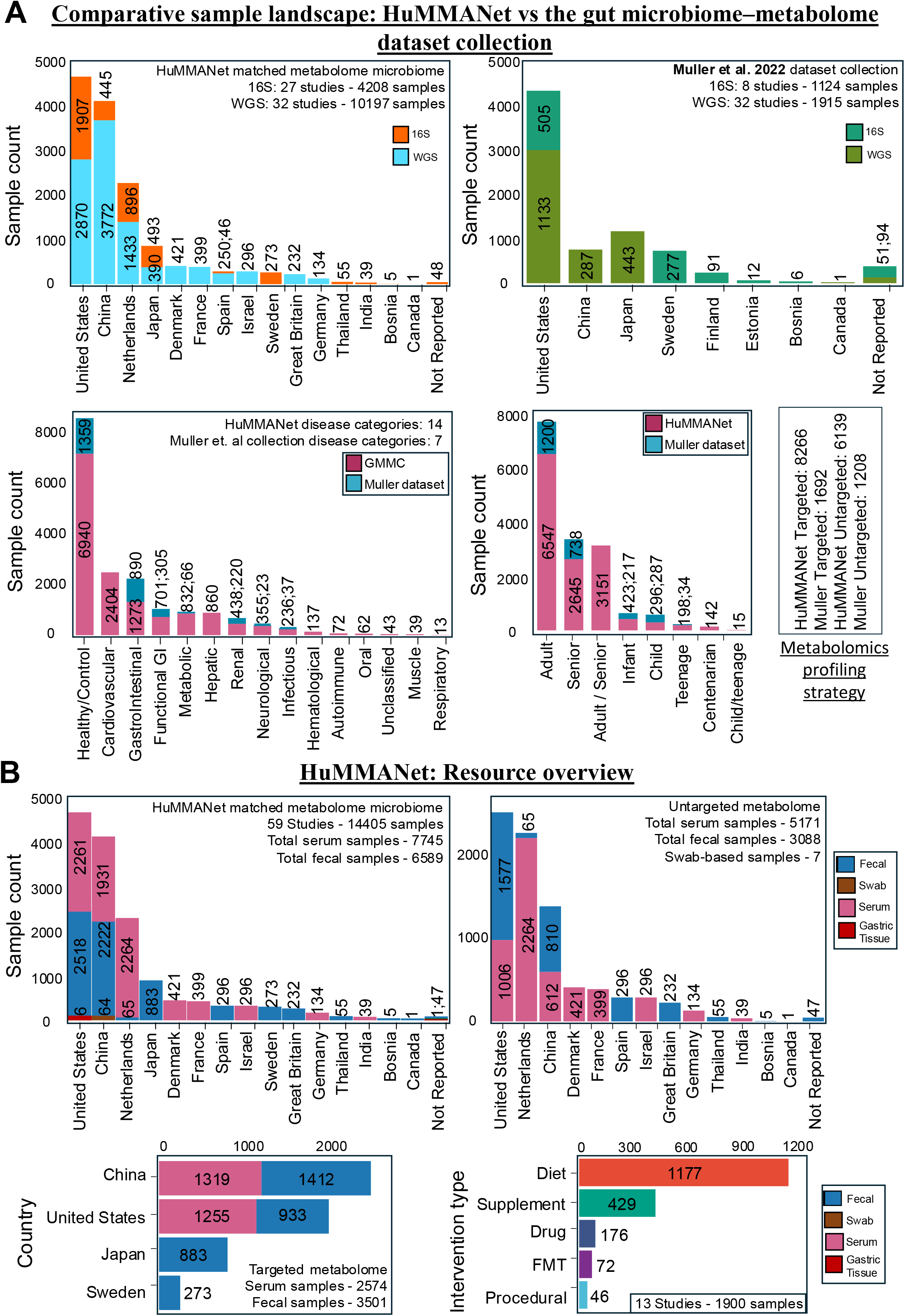
Comparison of HuMMANet and The Gut Microbiome–Metabolome Dataset Collection (Muller *et al* 2022) **(A)** Comparison of sample distribution across countries, disease categories, and age groups between the HuMMANet and the Muller et al. (2022) dataset collection^22^. HuMMANet includes 14,405 paired microbiome–metabolome samples, spanning both whole-genome shotgun (WGS) and 16S sequencing, with broader representation across disease categories and demographic groups. Differences in metabolomics coverage, including targeted and untargeted datasets, are also highlighted. **(B)** Sample distribution within HuMMANet by geography, biospecimen type (serum, fecal, swab, and gastric tissue), and metabolomics strategy (targeted and untargeted). Additional panels summarize country-wise contributions for targeted metabolomics and the distribution of intervention types (dietary, supplementation, pharmacological, fecal microbiota transplantation, and procedural studies; n = 12 interventional datasets, 1,815 samples).

Beyond observational cohorts, HuMMANet additionally incorporates 12 interventional datasets (1,815 samples) spanning dietary, supplementation, pharmacological, and fecal microbiota transplantation studies **(Figure 2B)**. HuMMANet also provides broad demographic and disease representation across age groups and clinical conditions (**Figure S1**). Together, these features establish HuMMANet as a comprehensive multi-omics resource for cross-study microbiome–metabolome analyses. Realizing this potential, however, first required resolving how the same metabolite is named across studies.

### Challenge of cross-study metabolite annotation variations

Despite harmonization of metadata and analytical workflows, cross-study metabolomics integration remained substantially constrained by inconsistent metabolite nomenclature and annotation. To quantify this limitation, we examined metabolite overlap as studies were progressively integrated. Individual studies reported 14 to 4,876 metabolites each, resulting in a total of 33,469 study-specific metabolite name entries **(Table S3)**, with 5,362 metabolites detected in at least two study cohorts and only three shared across 19 or more studies (out of the 59 study-cohorts) **(Figure 3A, left panel).**

**Figure 3:**
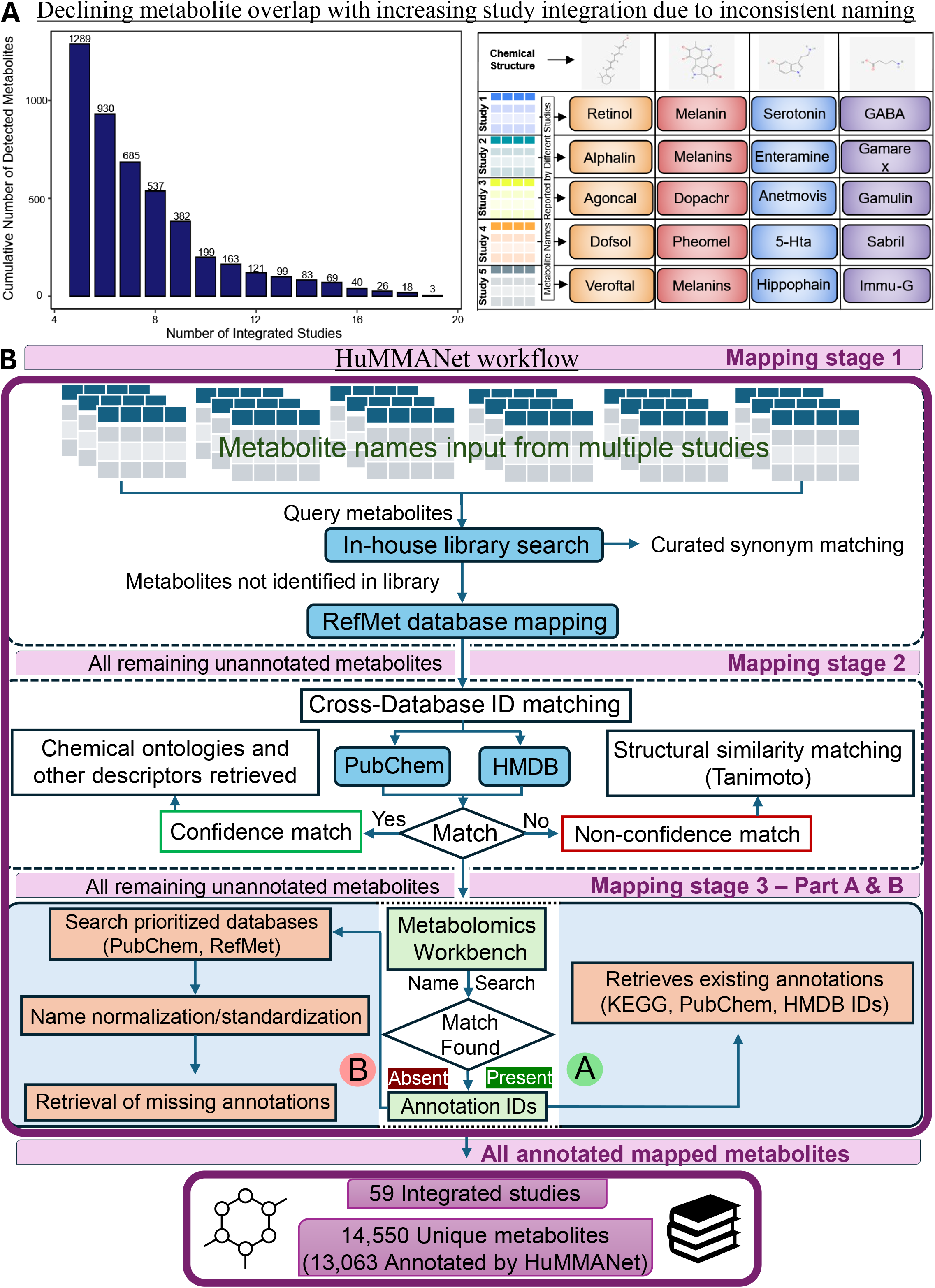
The need for cross-study metabolite harmonization, and the resulting development of the HuMMANet harmonization workflow. **(A)** Cross-study metabolomics integration is hindered by inconsistent metabolite naming across studies. As additional studies are integrated, the number of shared metabolites declines because identical chemical entities are reported under different names, reducing overlap despite representing the same molecular structures. **(B)** Overview of the HuMMANet workflow for harmonizing metabolite annotations across studies. The harmonization workflow proceeds in three stages. First, metabolite names from multiple datasets are standardized using RefMet (Stage 1) and existing annotation identifiers (PubChem, HMDB, ChEBI, KEGG) (Stage 2). Cross-database identifier matching identifies high-confidence matches, while conflicting annotations are retained with database provenance and evaluated using structural similarity. Remaining unmapped metabolites are resolved through prioritized offline PubChem searches and Metabolomics Workbench queries (Stage 3). The detailed explanation of the methodological steps in each stage is provided in **Text S1** and Methods section ‘Multi-stage metabolite annotation computational pipeline – HuMMANet workflow’

Inspection of the underlying annotations revealed the source of this fragmentation. Annotations varied widely — free-text names, database identifiers (HMDB, KEGG, PubChem), abbreviated synonyms, platform-specific labels, and unresolved m/z or retention-time features ^54,55^ — preventing direct cross-study matching (**Figure 3A, right panel; Table S2**). Consequently, identical metabolites were often represented by multiple annotations, fragmenting detection and limiting cross-study integration, meta-analysis, and identification of reproducible associations. These limitations of conventional identifier-based matching motivated the development of a multi-stage harmonization framework within HuMMANet that resolves annotation inconsistencies, recovers missing metabolite identities, and enriches harmonized metabolites with standardized biological annotations^29,39^.

### The core architecture of the HuMMANet harmonization pipeline

Having quantified the scale of the annotation problem, we next developed a framework to resolve it. This pipeline harmonized heterogeneous metabolite names to a unified nomenclature through a three-stage workflow **(Figure 3B),** consolidating metabolites from all 59 study-units into a common metadata structure with unique HuMMANet identifiers.

In stage one, HuMMANet mapped study-specific metabolite names to RefMet to retrieve standardized names, synonyms, cross-referenced identifiers, and chemical ontologies^40^ **(Methods**; **Text S1)**. RefMet contains >700,000 metabolites with associated synonyms and chemical ontologies^40,56^. For the current set of 59 study-cohorts, this step resulted in the annotation harmonization of 19,117 metabolite names **(Table S4)**. Remaining unannotated metabolites were carried forward to the second stage.

In stage two, remaining metabolites were queried against PubChem and HMDB using synonym- and identifier-based matching^41,42^ **(Methods; Text S1)**. This reconciled 8,645 additional metabolite annotations through high-confidence cross-database matches (‘Confidence Matches’) **(Table S4)**. Ambiguous mappings, i.e., where cross-referenced identifiers disagreed between databases, were classified as non-confidence matches and excluded from downstream harmonization, but provided with structural similarity scores and ontology annotations **(Table S5)**. All remaining unannotated metabolites (not belonging to Confidence and Non-Confidence match lists) were then investigated in the third stage.

In stage three, name normalization and extended Metabolomics Workbench searches recovered 225 additional metabolites (**Table S4**)^43^. The remaining 2,355 metabolites — either RefMet-matched without further annotation, or unmatched at every prior stage — were searched against an in-house curated PubChem library, which recovered more annotations than the online search despite higher computational cost.

Collectively, the pipeline reconciled 33,469 study-specific annotations into 14,550 unique metabolites. Following identifier harmonization, metabolites were enriched with biological knowledge layers, including metabolic pathways, microbial producer information, physiological context, and pharmacological annotations derived from SMPDB, MiMeDB-2.0, HMDB, DrugBank, and DrugCentral Databases^24,42,44–46^ **(Figure 4)**. This unified framework enables downstream cross-cohort comparisons, metabolite-centric network analyses, and biologically informed interpretation of shared metabolite signatures.

**Figure 4.**
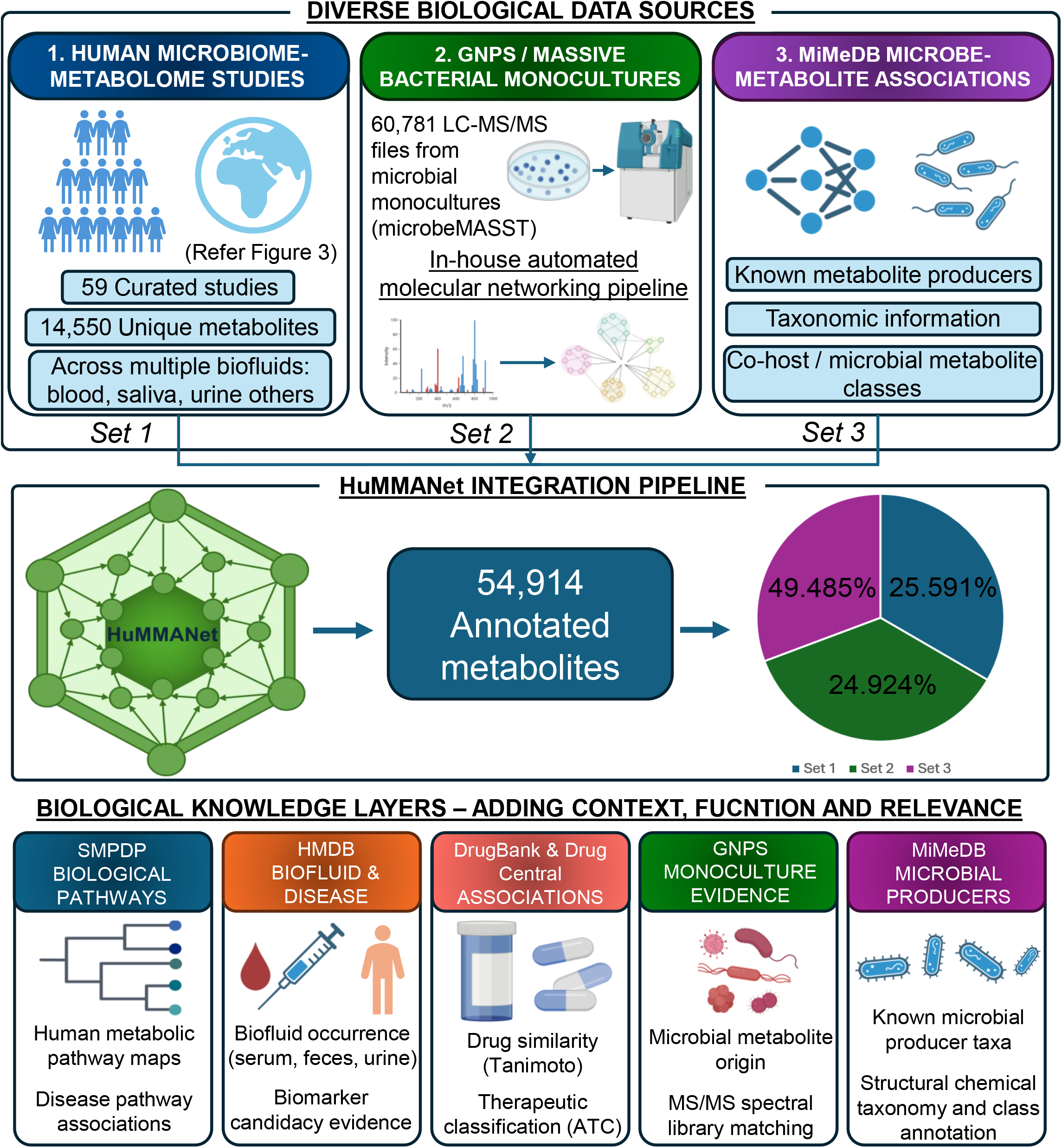
Construction and biological annotation of the HuMMANet metabolite atlas. The figure summarizes the construction of the HuMMANet metabolite atlas by integrating metabolites from three complementary biological data sources through the HuMMANet harmonization framework (**top panel**; Sets 1–3). **Set 1** comprises 14,550 unique metabolites from 59 curated human microbiome–metabolome study-units spanning multiple biofluids, whose annotation harmonization is described in Figure 3. **Set 2** comprises metabolite annotations derived from 60,781 LC–MS/MS files from bacterial monocultures (microbeMASST/GNPS-MassIVE), processed using an in-house automated molecular-networking pipeline to identify experimentally supported microbial metabolite associations. **Set 3** comprises curated microbe–metabolite associations from MiMeDB-2.0, including known microbial producers, taxonomic information, and host/microbial metabolite classifications. Metabolite annotations from Sets 1–3 were integrated using the harmonization workflow described in Figure 3, yielding a reference atlas of 54,914 unique metabolites (**middle panel**). The pie chart summarizes their source attribution: Set 1, 25.591%; Set 2, 24.924%; and Set 3, 49.485%. The harmonized atlas was subsequently enriched with complementary layers of biological and chemical context (**bottom panel**). These include SMPDB-derived pathway and disease-pathway annotations (7,232 metabolites linked to 48,642 pathway records); HMDB-derived biofluid occurrence and disease-biomarker evidence (6,073 metabolites linked to 2,253 biofluid records and 3,464 disease associations); DrugBank- and DrugCentral-derived structural similarity to approved drugs, assessed using Tanimoto coefficients against 4,947 and 4,099 drugs, respectively, together with therapeutic (ATC) classifications; GNPS monoculture-derived evidence of microbial metabolite origin; and MiMeDB-2.0-curated microbial producer information, complemented by structural chemical taxonomy and class annotations. Together, these layers extend HuMMANet beyond metabolite harmonization to establish a biologically contextualized reference atlas connecting harmonized metabolite identities with their microbial origins, biochemical pathways, physiological distributions, disease associations, chemical properties, and therapeutic relevance.

### Construction of the HuMMANet metabolite knowledgebase

Having resolved metabolite identities, we next asked what biological context could be attached to them. Each harmonized metabolite was annotated with standardized names, molecular formula, SMILES notation, and cross-reference identifiers to HMDB, PubChem, KEGG, ChEBI, and InChIKey, providing a common framework for cross-study integration and biological interpretation^41,42,57–59^. The resulting knowledgebase spanned amino acid derivatives, bile acids, lipids, organic acids, nucleotides, vitamins, neuroactive metabolites, and xenobiotic-associated compounds, each assigned a hierarchical chemical ontology (Super-Class, Main-Class, Sub-Class) enabling functional classification and enrichment analyses. Full annotation provenance was recorded for every metabolite, original study-specific name (Query Name), annotation source, identifier type/value, and originating study (Study Folder), allowing each metabolite to be traced back to its source.

Each metabolite was further linked to functional, physiological, microbial, and pharmacological context: 7,232 metabolites to 48,642 SMPDB pathway records via HMDB/KEGG/ChEBI identifiers^44^; 6,073 to 2,253 HMDB biofluid occurrence records (serum, feces, urine, saliva, CSF, sweat) and 3,464 disease associations across 523 phenotypes. Microbial context came from curated MiMeDB-2.0 metabolite–microbe associations (incorporating CMMC- KB) and experimentally derived producer annotations from GNPS/microbeMASST (covered in detail below)^24,26,47^. Pharmacological annotation compared metabolites against 4,947 DrugBank and 4,099 DrugCentral drugs using ECFP4 fingerprints (radius=2, 2048 bits; RDKit), ranked by Tanimoto similarity, retaining the top five matches per metabolite (high-confidence: Tanimoto ≥0.75)^45,46,60^.

Together, these layers transform HuMMANet from a harmonization framework into a comprehensive microbiome–metabolome knowledgebase.

### Integration of microbial monoculture metabolomics enables experimental annotation of putative microbial metabolite producers

Curated annotations alone, however, capture only associations that have already been reported. To add experimentally observed producer evidence, we next integrated microbeMASST (>60,000 microbial monoculture LC–MS/MS datasets spanning bacterial, archaeal, and eukaryotic taxa) ^26,47^, harmonizing GNPS molecular networking-derived metabolite features through the HuMMANet framework before integration^26,47^.

Across 60,781 LC–MS/MS files from microbial monoculture experiments, 198,402 metabolite features were detected, of which 156,541 (78.9%) were successfully harmonized, generating 196,352 unique microbe–metabolite associations linking 33,056 metabolites to 1,070 microbial species. Metabolite provenance analysis identified 149,807 metabolites detected exclusively in microbial monocultures and 48,595 shared with human biofluids (**Text S1; Figure S2**). These shared metabolites had 41,620, 24,923, 21,009, 10,288, 8,314, and 2,527 metabolite– biofluid occurrences in serum, urine, feces, saliva, cerebrospinal fluid, and sweat, respectively.

To facilitate future expansion, we developed an automated GNPS molecular networking pipeline for high-throughput processing of microbial monoculture datasets (**Text S2**). The resulting harmonized microbe–metabolite associations are integrated into HuMMANet and distributed through the accompanying GitHub repository, providing a reference framework for assigning putative microbial producers to metabolites.

### Expanding biochemical coverage through integration of the Microbial Metabolome Database

Having incorporated experimentally derived producer annotations, we next sought to extend biochemical coverage beyond the metabolites identified in HuMMANet study cohorts, and integrated the 2026 release of MiMeDB (MiMeDB-2.0)^24^, which also includes curated compound entries from the CMMC-KB^25^, into the HuMMANet knowledgebase (**Figure 4**)^24^. Following identifier harmonization, synonym reconciliation, chemical ontology mapping, and integration with study-derived metabolites, the resulting knowledgebase comprised 54,914 unique metabolites. This standardized reference enables direct mapping of future microbiome– metabolome datasets onto a common annotation space, with only metabolite annotations that cannot be matched to the existing reference requiring harmonization. Together, harmonized study-derived and curated microbial metabolites provide a unified reference for cross-study integration and biological interpretation.

### Comparative mapping performance of HuMMANet

With the resource assembled, we next asked how it performs relative to existing tools. To systematically position HuMMANet relative to existing approaches, we compared its functional capabilities with widely used metabolite annotation and mapping frameworks **(Table 1)**. Most existing methods, including MetaboAnalyst, IDmapping, and MetaboliteAnnotator, annotate metabolites within individual datasets without supporting cross-study harmonization^29–31^. Model-centric tools like MetaboAnnotator instead map metabolites onto VMH/genome-scale reconstructions, a distinct paradigm not built for cross-study integration^61^. On the other hand, while metID is LC–MS-specific^32^, MAPS requires pre-generated outputs from MZmine, GNPS2, SIRIUS, and MS2Query^34–38,62^. These limit their applicability where such intermediate outputs are unavailable. HuMMANet, by contrast, annotates and harmonizes across studies in a single workflow independent of external pipeline outputs, retaining cross-references at the user’s preferred identifier level.

**Table 1.**
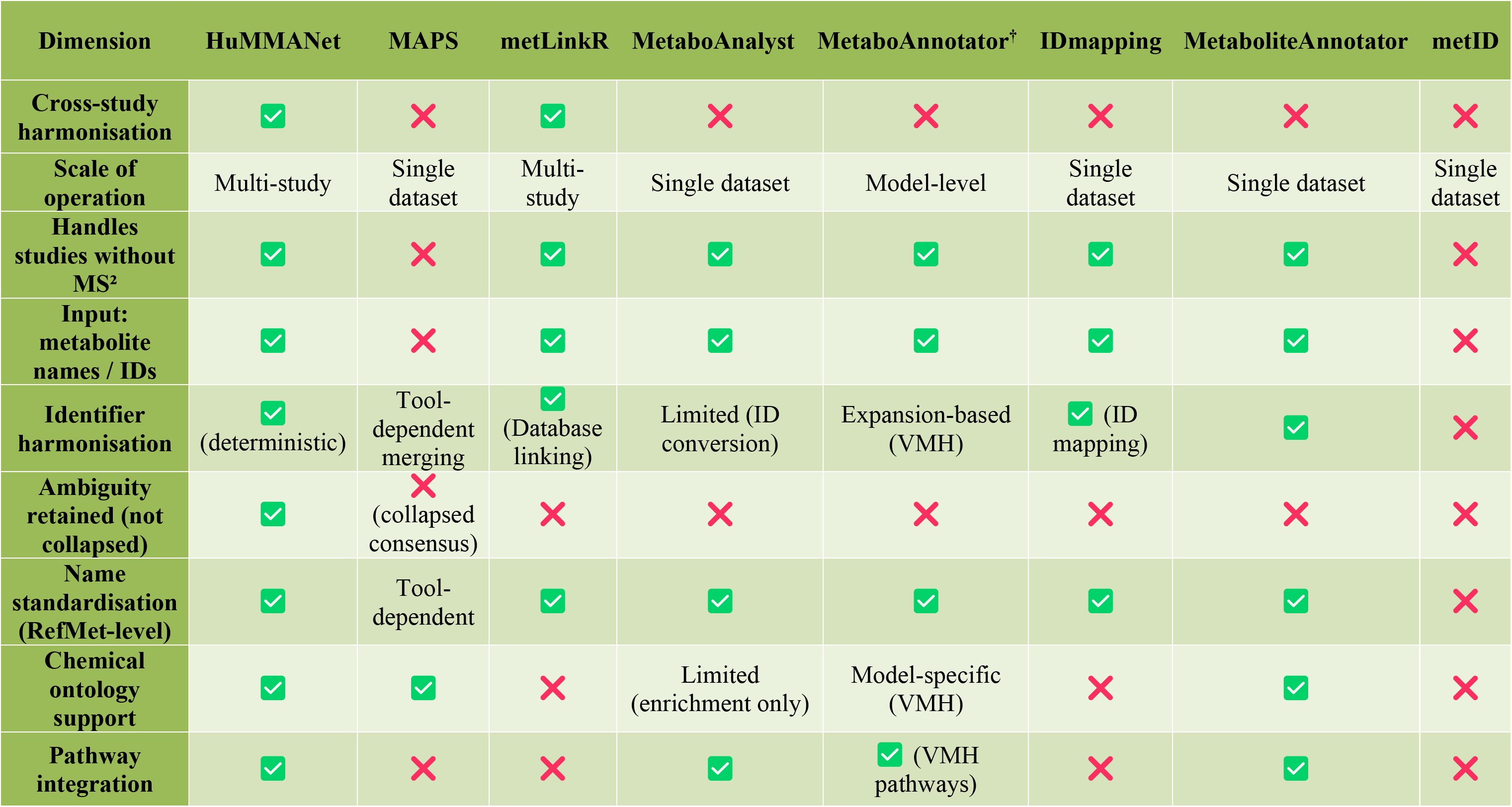
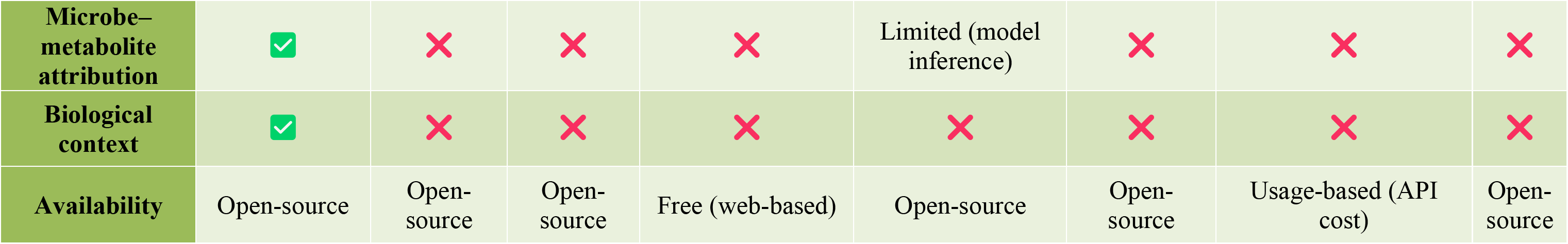
Functional comparison of metabolite annotation and mapping frameworks. Comparison of HuMMANet with commonly used tools across key functional dimensions, including cross-study harmonization, scale of operation, identifier standardization, ambiguity handling, and integration of biological context. Checkmarks (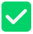) indicate a supported feature and crosses (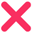) indicate an unsupported one; for dimensions where support is partial, conditional, or implemented through a different mechanism (e.g., Scale of operation, Identifier harmonisation, Chemical ontology support, Pathway integration, Microbe–metabolite attribution, Availability), the specific implementation is noted in place of a binary symbol. HuMMANet uniquely supports deterministic cross-study harmonization with preservation of annotation ambiguity and integrated biological interpretation layers, including chemical ontology, pathway mapping, and microbe–metabolite attribution. Other methods provide partial functionality, typically restricted to single-dataset processing, identifier conversion, or model-specific annotation. †MetaboAnnotator is a model- centric tool that maps metabolite names and identifiers onto genome-scale metabolic reconstructions and VMH nomenclature rather than performing cross-study metabolomics harmonization, and is included here for functional comparison only; it was not included in the empirical benchmarking shown in **Figure 5A** (benchmarked against MetaboAnalyst, MetaboliteAnnotator, IDmapping, and metLinkR). Abbreviations: MS, mass spectrometry; RefMet, reference metabolite nomenclature standard; VMH, Virtual Metabolic Human.

Among the frameworks evaluated here, metLinkR is the only tool that similarly performs multi-study annotation^39^. However, metLinkR standardizes metabolite names without integrating complementary chemical descriptors, ontology annotations, or cross-referenced database identifiers that facilitate downstream biological interpretation and cross-resource interoperability. To assess HuMMANet’s annotation performance, we compared it with MetaboAnalyst, MetaboliteAnnotator, IDmapping, and metLinkR **(Figure 5A; Tables S6-S7)**. Benchmarking against six MetaboLights datasets (MTBLS11733, MTBLS12636, MTBLS13039, MTBLS12997, MTBLS13105, and MTBLS12764), previously used to evaluate MetaboliteAnnotator, was performed using its non-AI workflow to ensure transparent and reproducible comparisons^31,63^. Annotation performance, the proportion of input metabolite names mapped to standardized identifiers, was highest for HuMMANet across all six datasets in both positive- and negative-ionization modes **(Figure 5A, left panels)**. For comparison with metLinkR, we used the same five benchmarking datasets (Broad 2022, Vicky 2019, DDLPS, LECOC, and COMETS) originally employed in the metLinkR evaluation^39^**(Figure 5A; right panel)**. HuMMANet outperformed metLinkR across all datasets, with comparable performance only for Vicky 2019.

**Figure 5:**
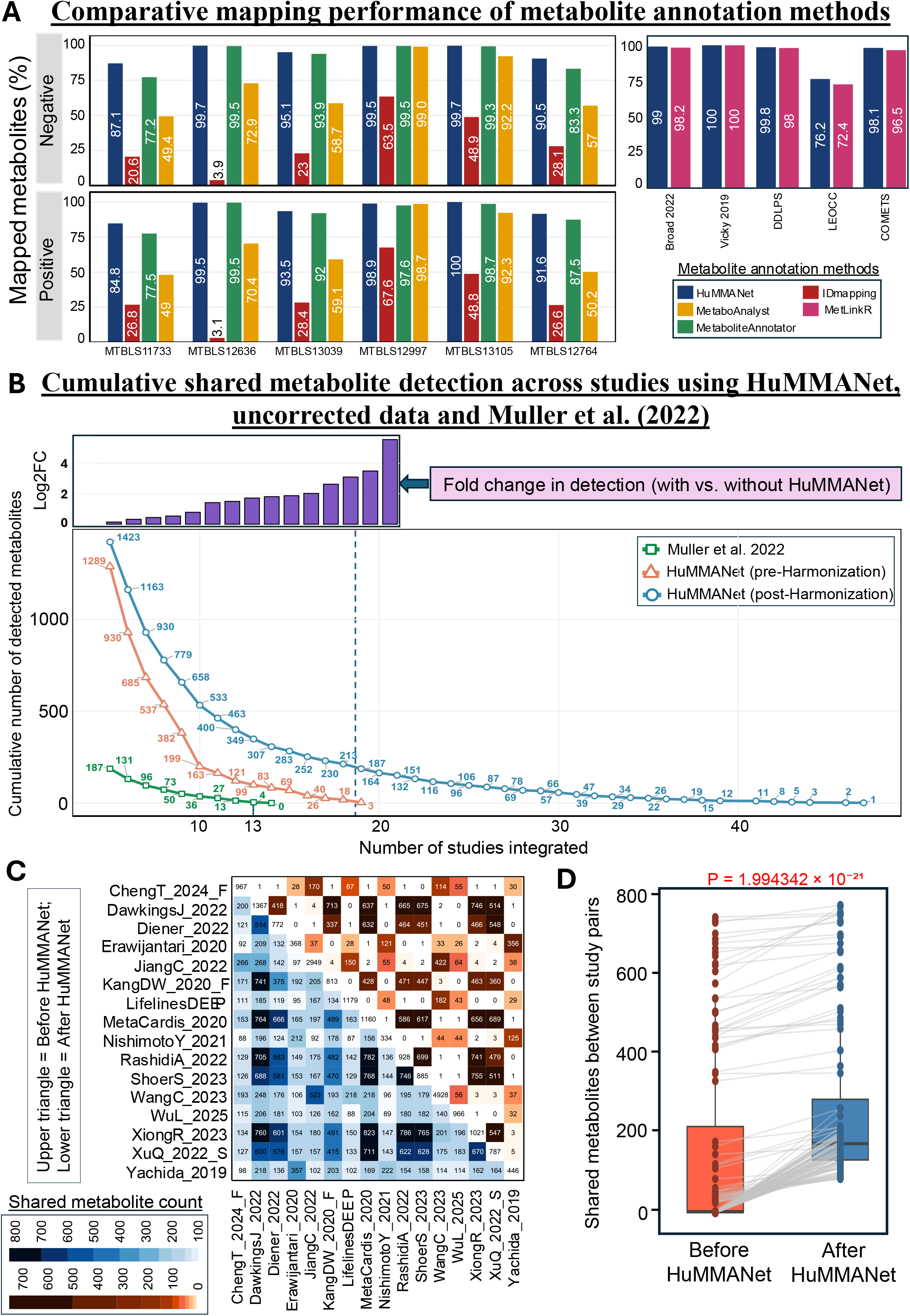
HuMMANet Expands Cross-Study Metabolite Harmonization and Outperforms Existing Annotation Approaches. **(A) Left panel**: Percentage of mapped metabolites across multiple studies, stratified into negative and positive ionization modes, comparing HuMMANet with existing annotation tools (MetaboAnalyst, MetaboliteAnnotator, IDmapping). The six MetaboLights datasets were selected because they were previously used to benchmark MetaboliteAnnotator, enabling direct comparison with established annotation tools. **Right panel**: Percentage of mapped metabolites comparing HuMMANet with metLinkR across external benchmark datasets. The five datasets were selected because they were used in the original metLinkR evaluation, enabling direct comparison with a multi-study annotation framework. **(B)** The cumulative number of detected metabolites is shown as a function of the number of studies integrated, comparing with HuMMANet (post-harmonization), metabolite profiles (without harmonization), and the Muller et al. (2022) dataset collection^22^. HuMMANet consistently increases metabolite detection across all integration levels, resulting in an expanded shared metabolite space relative to both baseline and prior approaches. The inset shows the fold change in metabolite detection (log₂ scale) for HuMMANet relative to the baseline (with vs without HuMMANet), demonstrating progressive gains in detection as more datasets are integrated. **(C)** Pairwise comparison of shared metabolites across studies in HuMMANet, shown as a symmetric heatmap. The upper triangle represents shared metabolite counts prior to HuMMANet harmonization, while the lower triangle represents shared counts after harmonization. HuMMANet substantially increases cross-study metabolite overlap, expanding the shared metabolite space and improving comparability across datasets. **(D)** Quantitative summary of the pairwise shared metabolite counts shown in panel C. Each point pair represents the number of metabolites shared between the same pair of studies before and after HuMMANet harmonization, with grey connecting lines indicating paired comparisons. Boxplots summarize the distribution of shared metabolite counts across all study pairs. HuMMANet significantly increases cross-study metabolite overlap (paired Wilcoxon signed-rank test, P = 1.99 × 10⁻²¹), demonstrating a systematic expansion of the harmonized metabolite space across the HuMMANet collection

We next evaluated the impact of harmonization on metabolite sharing across progressively integrated cohorts (**Figure 5B**). Without harmonization, shared metabolites declined rapidly, with none detected across more than 19 studies, consistent with the Curated Gut Microbiome– Metabolome collection. HuMMANet, with its harmonization step, substantially increased shared metabolite retention at every integration depth, maintaining overlap beyond 45 studies and achieving up to 62-fold higher shared metabolite detection at the 19-study threshold (**Figure 5B**, inset).

To assess study-level integration, we compared shared metabolites across all 1,711 study pairs before and after harmonization (listed in **Table S8**; with comparison data shown for a subset in **Figure 5C**). HuMMANet significantly increased metabolite overlap (paired Wilcoxon signed-rank test, P = 1.99 × 10⁻²¹) (**Figure 5D**). Among the 583 of the 1,711 study-pairs (34.1%) with no shared metabolites prior to harmonization, a median of 21 shared metabolites was recovered per pair. For the remaining study pairs, harmonization increased shared metabolites by a mean of 19- fold, with a maximum improvement of 218-fold (**Table S8**). Together, these benchmarks show that harmonization substantially expands the metabolite space available for cross-study analysis, a gain we next exploited biologically.

### HuMMANet facilitates identification of consistent microbiome-associated metabolites with specific chemical and functional characteristics

Having established that harmonization expands cross-study metabolite coverage, we next asked whether this gain translates into biological insight. To demonstrate HuMMANet’s utility, we applied it to paired adult gut microbiome–metabolome profiles: 7,603 microbiomes across 19 cohorts with serum metabolome data, and 4,813 microbiomes across 30 cohorts with fecal metabolome data to identify metabolites whose levels showed consistently significant associations with gut microbiome composition across datasets (**Methods**; **Figure S3**). Examining shared metabolite detection as a function of the minimum number of studies in which each metabolite was detected (study threshold; **Figure 6A**) showed that the application of the HuMMANet framework substantially improved cross-study metabolite retention in both metabolomes, increasing progressively as more studies were integrated (**Figure S4**). Associations between retained metabolites and gut microbial composition were evaluated independently per study using PERMANOVA; metabolites significantly associated (P < 0.05) in at least 50% of the studies in which they were detected were classified as microbiome-associated. Of the 4,267 unique serum metabolites detected in at least one study, only 519 (∼12%) showed significant microbiome associations at this minimal detection threshold. This proportion rose sharply as the reproducibility requirement increased, reaching 57% at a ≥5-study threshold, peaking at 59% at ≥6 studies, and settling between 40–46% at the more stringent ≥10–12-study thresholds (**Figure 6A**, **left panel**). Similarly, while only 322 of the 7,962 unique metabolites detected in at least one paired adult fecal cohort (∼4%) were significantly associated with gut microbiome composition, this proportion increased markedly with cross-study reproducibility, exceeding 50% across the ≥5–10 study thresholds (**Figure 6A**, **right panel**). Thus, filtering by increasing cross-study reproducibility, enabled by HuMMANet’s harmonization workflow, enriches for biologically meaningful, microbiome-linked metabolites across both metabolome types.

**Figure 6.**
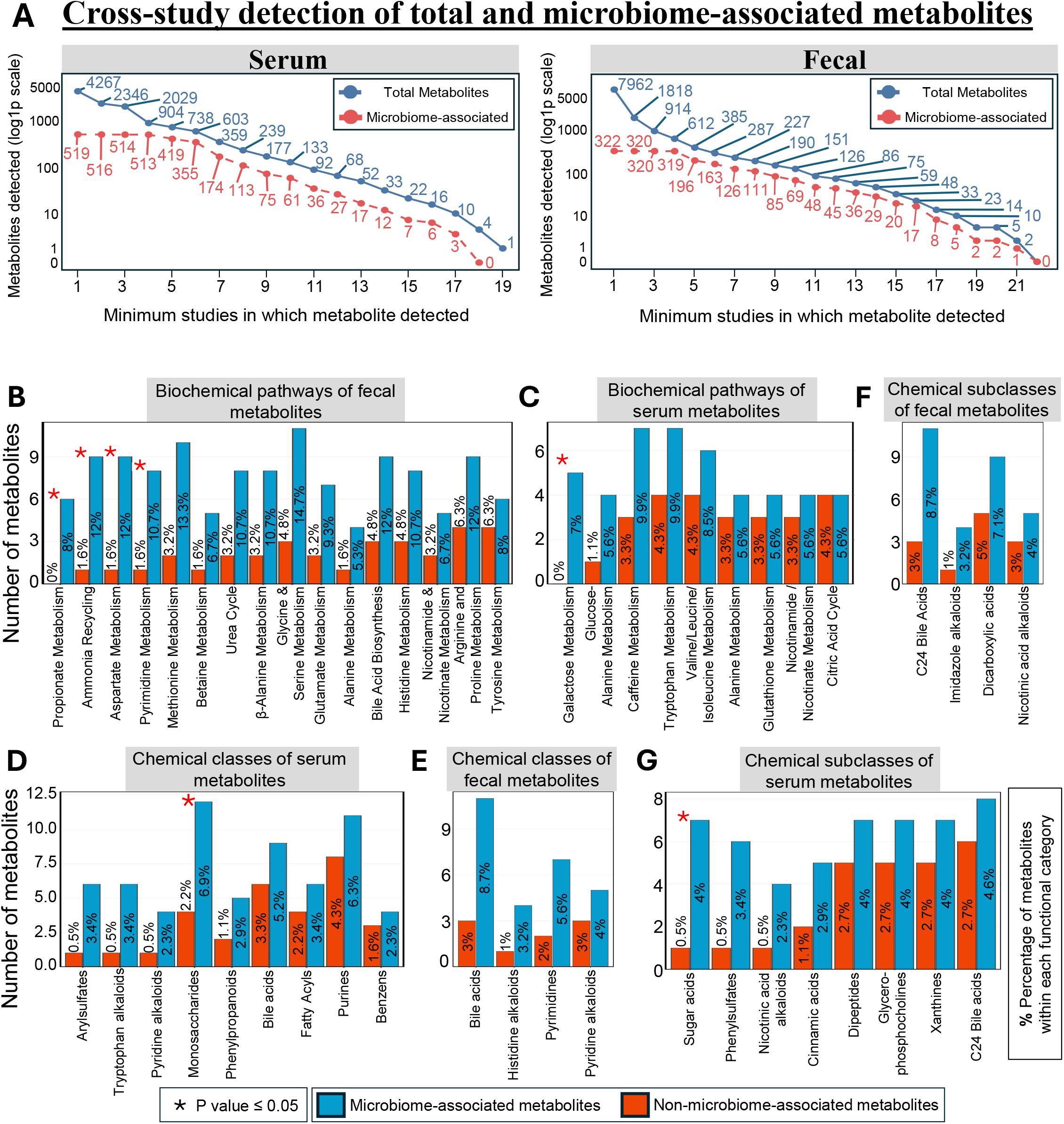
Cross-study reproducibility identifies microbiome-associated metabolites with conserved pathway and chemical class enrichment. (**A**) Cumulative numbers of total metabolites (blue) and microbiome-associated metabolites (red) identified across increasing minimum study-occurrence thresholds for serum (left) and fecal (right) metabolomes. At each threshold, blue lines indicate the number of metabolites detected in at least the specified number of studies. Red lines indicate the subset of these metabolites that were significantly associated with gut microbial community composition (PERMANOVA, P ≤ 0.05) in at least 50% of the studies in which they were detected. Numbers indicate metabolite counts at each study-occurrence threshold. **B-C.** Enrichment of microbiome-associated metabolites across biochemical pathways in fecal (**B**) and serum (**C**) metabolomes. Blue bars represent microbiome-associated metabolites and orange bars represent non-microbiome-associated metabolites. Numbers within bars indicate the percentage of metabolites within each pathway classified as microbiome-associated or non-microbiome-associated. Asterisks denote pathways significantly enriched for microbiome-associated metabolites (Fisher’s exact test, *P* ≤ 0.05). **D-E.** Enrichment of microbiome-associated metabolites across major chemical classes in serum (**D**) and fecal (**E**) metabolomes. Blue bars represent microbiome-associated metabolites and orange bars represent non-microbiome-associated metabolites. Percentages indicate the proportion of metabolites assigned to each chemical class. Asterisks indicate significant enrichment (Fisher’s exact test, *P* ≤ 0.05). **F-G.** Enrichment of microbiome-associated metabolites across chemical subclasses in fecal (**F**) and serum (**G**) metabolomes. Blue and orange bars represent microbiome-associated and non-microbiome-associated metabolites, respectively, with percentages indicating the fraction of metabolites assigned to each subclass. Asterisks denote significantly enriched subclasses (Fisher’s exact test, *P* ≤ 0.05). Complete enrichment statistics are provided in **Tables S9–S14**.

We adopted a cross-study threshold of ≥7 studies for downstream analyses, balancing reproducibility with metabolite retention. At this threshold, 48.5% (174/359) of retained serum metabolites and 55.5% (126/227) of retained fecal metabolites were classified as microbiome-associated and were investigated further.

We next assessed whether these metabolites showed distinct functional signatures, comparing their distribution across metabolic pathways and chemical ontologies against non-associated metabolites using fold enrichment and Fisher’s exact tests (**Figures 6B-6G**; **Tables S9- S14**). In feces, propionate metabolism, ammonia recycling, pyrimidine metabolism, and aspartate metabolism were significantly enriched (**Figure 6B**; **Table S9**); methionine, glycine/serine, histidine, glutamate, and tyrosine metabolism, the urea cycle, betaine metabolism, and nicotinate/nicotinamide metabolism showed high but non-significant enrichment. Betaine, notably, derives from dietary choline and feeds gut microbial trimethylamine production, linking this pathway to an established host–microbiome axis^64,65^. In serum, galactose metabolism was the only significantly enriched pathway (**Figure 6C**; **Table S10**); tryptophan metabolism showed a strong non-significant trend. This was notable because tryptophan-derived metabolites act as microbiota-derived signals regulating host immunity and gut–brain communication^66,67^.

Similarly, chemical-class analysis showed similar concentration within specific families: monosaccharides were the only significantly enriched serum class (**Figure 6D**; **Table S11**), while no fecal class reached significance (**Figure 6E; Table S12**). At the subclass level, C24 bile acids, imidazole alkaloids, and dicarboxylic acids were significantly enriched in feces (**Figure 6F; Table S13**), consistent with gut bacteria’s established role converting primary to secondary bile acids that regulate host metabolism and immune signaling^68,69^. In serum, sugar acids were significantly enriched (**Figure 6G; Table S14**), consistent with carbohydrate-derived metabolites’ role in host– microbiome interactions^70^. In addition, phenylsulfates, nicotinic acid alkaloids, cinnamic acids, dipeptides, glycerophosphocholines, xanthines, and C24 bile acid derivatives showed similar non-significant trends. Together, microbiome-associated metabolites preferentially cluster within specific pathways and chemical families rather than distributing uniformly across the metabolome.

### HuMMANet identifies metabolite panels positively and negatively associated with a health-associated resilient microbiome

We next asked whether microbiome-associated metabolites preferentially align with health-linked taxa. A previous study from our group investigated 45,424 adult human gut microbiomes worldwide, ranking 201 taxa by the consistency of their positive associations with three key properties: prevalence/community association in non-diseased subjects, longitudinal stability, and host health. The resulting score was termed the HACK index, where taxa higher in this ranking more consistently associate with a resilient, health-associated microbiome^48^. To identify metabolites reflecting this resilient state, we leveraged this framework. Of the HACK-ranked taxa investigated in the original study^48^, 196 overlapped with HuMMANet’s taxa abundance profile and were therefore included in the analysis. Metabolites whose taxon-level association patterns positively correlate with taxon-specific HACK scores (more positive associations with high- HACK taxa, more negative with low-HACK taxa) likely mark a health-associated, stable microbiome, while those showing the reverse trend likely mark detrimental configurations. Our objective was to identify metabolite panels capable of distinguishing between these two configurations.

Focusing on metabolites detected in ≥7 studies, and analyzing serum and fecal datasets separately, we computed an Association-Score between each metabolite and each HACK-ranked taxon^71^ (**Text S3**; **Figure S3**). This score captures both the direction and cross-cohort consistency of the metabolite-taxon relationship: its sign indicates direction, and its magnitude reflects consistency across cohorts. For each metabolite, we correlated its Association-Score profile across the 196 taxa with their corresponding HACK scores. Metabolites showing positive correlations, preferentially associated with consistent, health-linked core-keystones, were termed HACK- positive, while those showing negative correlations, preferentially associated with dysfunction-linked taxa, were termed HACK-negative. This identified 58 serum and 25 fecal HACK-positive metabolites, and 38 serum and 65 fecal HACK-negative metabolites (**Table S15**). These associations were robust to the inclusion of studies from the original HACK discovery cohort: a sensitivity analysis excluding them yielded highly concordant scores for both fecal (R = 0.89, P < 2.2×10⁻¹⁶) and serum (R = 0.61, P < 5.7×10⁻³³) metabolites (**Figure S5**).

Having defined these panels, we next examined the individual metabolites driving them. In serum, the two metabolites most strongly positively associated with taxon-level HACK indices were 1H-indole-3-propanoic acid (IPA; R=0.5, p=8.6×10⁻¹⁴) and 3-phenylpropionate (R=0.45, p=4.4×10⁻¹¹) (**Figure 7A**), highlighting circulating tryptophan catabolites and phenolic acid metabolites as characteristic outputs of health-associated gut commensals. IPA is a gut microbiota-derived tryptophan metabolite that activates the pregnane X receptor (PXR), suppresses TLR4- mediated inflammatory signaling, and supports intestinal tight-junction integrity; reduced circulating IPA is consistently reported in metabolic and inflammatory disorders^72^. Supplementation of IPA has also been associated with increased responsiveness to immune-checkpoint blockade therapy in multiple cancers^73^. Because IPA is produced by gut bacteria from dietary tryptophan, its recovery here is a particularly compelling demonstration of HuMMANet’s ability to retrieve microbiome-associated circulating signals^74^. 3-Phenylpropionate, another microbial metabolite arising from aromatic-compound and polyphenol metabolism, showed a similarly strong positive association with HACK scores, further supporting its link to health-associated microbial communities^75,76^.

**Figure 7:**
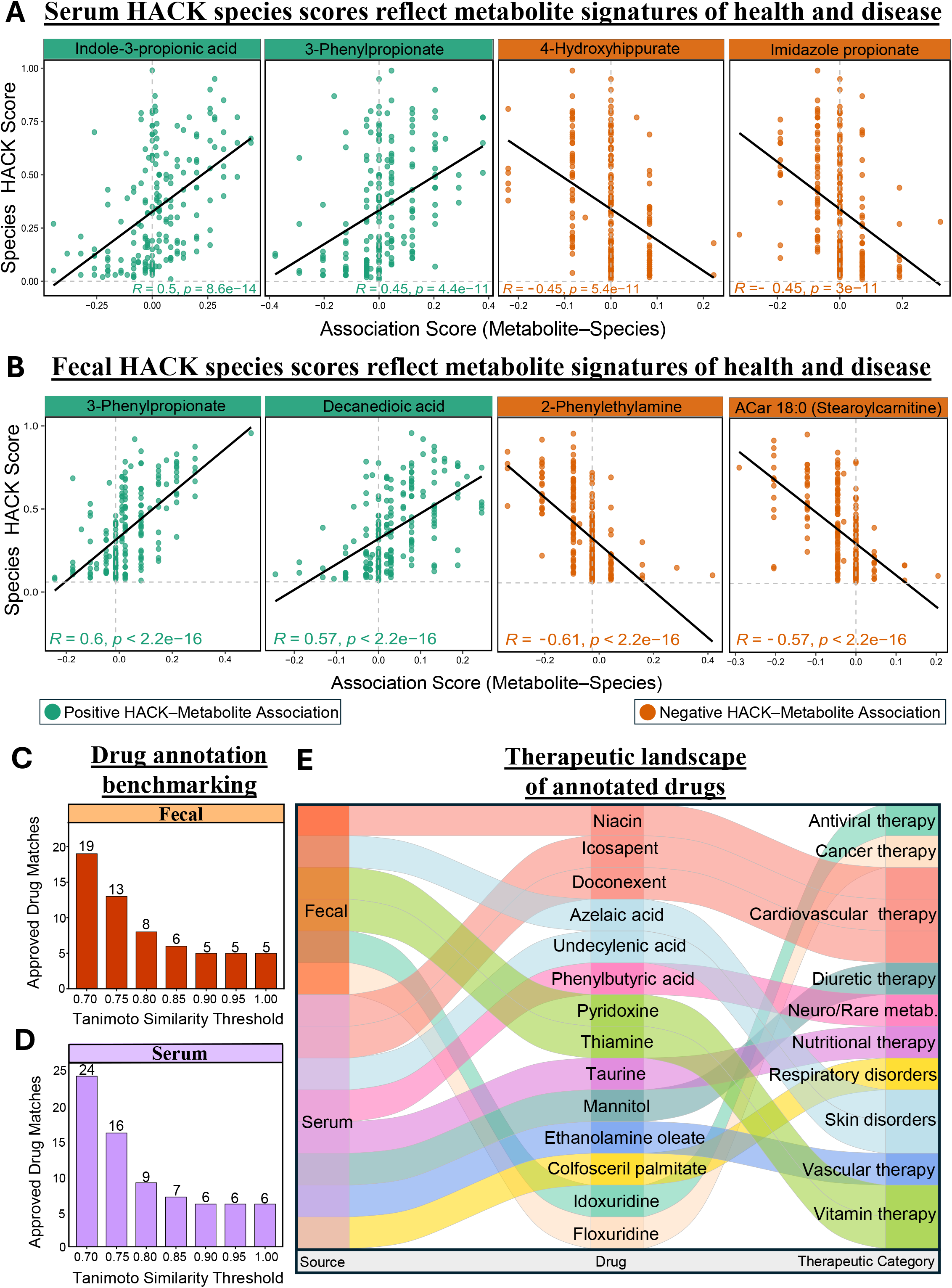
HACK-associated metabolites reveal conserved signatures of microbiome health and structural similarity to clinically used drugs. **(A)** Representative serum metabolites showing the strongest positive (green; indole-3-propionic acid, R=0.5, P=8.6×10⁻¹⁴; 3-phenylpropionate, R=0.45, P=4.4×10⁻¹¹) and negative (orange; 4- hydroxyhippurate, R=−0.45, P=5.4×10⁻¹¹; imidazole propionate, R=−0.45, P=3×10⁻¹¹) associations with taxon-specific Health-Associated Core Keystone (HACK) scores. Each point represents one HACK-ranked microbial taxon; lines show linear regression fits. **(B)** Representative fecal metabolites showing the strongest positive (green; 3-phenylpropionate, R=0.6, P<2.2×10⁻¹⁶; decanedioic acid, R=0.57, P<2.2×10⁻¹⁶) and negative (orange; 2-phenylethylamine, R=−0.61, P<2.2×10⁻¹⁶; ACar 18:0 [stearoylcarnitine], R=−0.57, P<2.2×10⁻¹⁶) associations with HACK scores. Positive correlations indicate preferential association with health-associated microbial taxa; negative correlations indicate preferential association with taxa linked to microbiome dysfunction**. C-D.** Number of unique approved drugs showing structural similarity to HACK- positive microbiome-associated metabolites across increasing Tanimoto similarity thresholds in fecal (**C**) and serum (**D**) metabolomes, after combining DrugBank and DrugCentral annotations (13 fecal and 16 serum metabolites reached high-confidence matches, Tanimoto ≥0.75). **(E)** Representative high-confidence structural analogues (Tanimoto similarity ≥ 0.75) linking HACK- positive microbiome-associated metabolites to approved therapeutics across diverse therapeutic categories. The Sankey diagram shows whether each metabolite–drug match originated from serum or fecal metabolites (left), the corresponding approved drug analogue (middle), and its primary therapeutic category (right). Representative drug analogues span cardiovascular, skin, neurological and rare metabolic, vitamin, nutritional, diuretic, vascular, respiratory, antiviral, and cancer therapies. Complete structural similarity results are provided in **Table S16**.

In contrast, the two most negatively associated serum metabolites were 4- hydroxyhippurate (R=−0.45, p=5.4×10⁻¹¹) and imidazole propionate (R=−0.45, p=3×10⁻¹¹). The latter is a well-established inhibitor of insulin signalling via mTORC1 activation, produced from histidine by dysbiosis-associated bacteria; elevated levels are consistently reported in type 2 diabetes and metabolic syndrome, and have more recently been linked to neuroinflammatory mechanisms^77–79^.

In fecal metabolomes, 3-phenylpropionate again showed the strongest positive association with HACK scores (R=0.6, p<2.2×10⁻¹⁶), together with decanedioic acid (R=0.57, p<2.2×10⁻¹⁶), consistent with microbial aromatic-compound metabolism and fatty-acid catabolism, respectively (**Figure 7B**)^80,81^. The two most negatively associated fecal metabolites were 2-phenylethylamine (R=−0.61, p<2.2×10⁻¹⁶) and ACar 18:0 (stearoylcarnitine; R=−0.57, p<2.2×10⁻¹⁶) (**Figure 7B**). 2- Phenylethylamine has previously been linked to altered aromatic amino acid metabolism in dysbiotic microbial communities^82^, and ACar 18:0 (stearoylcarnitine) is a long-chain acylcarnitine whose fecal elevation has been associated with altered intestinal epithelial mitochondrial fatty-acid metabolism in inflammatory bowel disease^83^. Together, these panels nominate specific metabolites as reproducible markers of microbiome health and dysbiosis, raising the question of their biological origin and translational relevance.

### Biological annotation links HACK-associated metabolites to clinically relevant drugs and putative microbial producers

Having identified metabolites that track with microbiome health, we finally asked whether they hold translational relevance. To explore the therapeutic relevance of HACK-positive metabolites, we assessed how closely they resembled clinically used drugs, comparing molecular structures against approved drugs from DrugBank and DrugCentral using Tanimoto structural similarity^45,46^, a feature already incorporated within HuMMANet. Thirteen fecal and 16 serum HACK-positive metabolites showed strong structural similarity (Tanimoto ≥ 0.75) to approved drugs (**Table S16**, **Figures 7C-D**), spanning categories including cardiovascular, neurological, rare metabolic, respiratory, nutritional, skin, vitamin, diuretic, vascular, antiviral, anti-inflammatory, and cancer therapies (**Figure 7E**). Among high-confidence matches, 3-phenylpropionate resembled phenylbutyric acid (Tanimoto = 0.81), used for urea cycle disorders and ALS; EPA and DHA resembled the drugs icosapent and doconexent; and dicarboxylic acids (decanedioic, octanedioic, undecanedioate, dodecanedioate) resembled azelaic acid, used for inflammatory skin disorders^84^.

Several HACK-positive vitamins (thiamine, pyridoxine, niacin, nicotinamide) showed exact matches (Tanimoto = 1.0) to approved compounds, consistent with gut microbes’ known ability to synthesize B vitamins^4^. Nucleosides adenosine, cytidine, and thymidine resembled the antiviral/anticancer drugs vidarabine, cytarabine, idoxuridine, and floxuridine (**Table S16**). These findings show that microbiome-associated metabolites share structural similarity with clinically used drugs across therapeutic areas; while structural similarity alone does not imply shared activity, it offers a basis for prioritizing metabolite panels for future study. To aid interpretation of these panels, we annotated experimentally reported microbial producers using the HuMMANet- integrated reference database, combining GNPS-derived monoculture annotations with curated MiMeDB-2.0 associations (**Text S3**; **Table S17**).

## Discussion

Cross-study microbiome–metabolome integration has long been constrained by inconsistent metabolite annotation, limiting reproducible comparisons across studies and obscuring conserved host–microbiome metabolic signatures^22,31,39^. Here, we address this gap with a curated gut microbiome–metabolome resource offering three advances over prior efforts. First, it provides a >4-fold expansion in paired samples (46 studies, 59 cohort-assay pairs, 14,405 samples, 13 disease categories plus healthy/control vs. 7 previously), with broader geographic and sequencing- modality coverage (16S + WGS; targeted + untargeted metabolomics). Second, its multi-stage harmonization framework reconciled 33,469 study-specific annotations into 14,550 harmonized metabolites. This set was expanded to 54,914 through integration with MiMeDB-2.0, which incorporates the CMMC-KB, a curated resource incorporating experimentally supported microbial metabolite annotations, including evidence from germ-free versus colonized animal models^25^, and further supplemented with >60,000 microbeMASST monoculture-derived metabolites^24,26^. Third, this repository is linked to a knowledgebase built by connecting these annotations to external resources including RefMet, HMDB, PubChem, Metabolomics Workbench, SMPDB, MiMeDB- 2.0, GNPS-microbeMASST, DrugBank, and DrugCentral, which supply standardized chemical identifiers, biochemical pathways, putative microbial producers, physiological distributions, disease links, and therapeutic structural relationships for each metabolite^24,26,40–47^.

Benchmarking against established annotation tools demonstrated that HuMMANet achieves superior annotation performance while uniquely combining multi-study harmonization with rich biological context, a capability no published tool provides. This translated into substantial practical gains, with shared-metabolite detection improved up to 62-fold at deeper integration thresholds and a mean 19-fold (up to 218-fold) increase in detection overlap in study pairs, including recovery of shared metabolites for over a third of pairs that had none in common prior to harmonization. By providing a standardized reference while preserving study-level information, HuMMANet improves interoperability across heterogeneous datasets and supplies a reusable foundation for reproducible analyses. This in turn facilitated the discovery of robust microbiome– metabolome links: metabolites consistently detected across multiple cohorts were markedly more likely to associate significantly with gut microbial community composition than those detected only sporadically, supporting cross-study reproducibility as a practical criterion for prioritizing biologically meaningful microbiome-associated metabolites. It was also evident in our identification of a robust panel of HACK-positive metabolites that can be prioritized for further mechanistic and translational evaluation.

The utility of this integrated annotation framework is further demonstrated in our proof-of-concept application, where we show that microbiome-associated metabolites are not randomly distributed throughout the metabolome but preferentially cluster within conserved host– microbiome metabolic processes, including amino acid, short-chain fatty acid, bile acid, carbohydrate, and nucleotide metabolism. Integration with the HACK framework extended these observations by identifying metabolites reproducibly associated with health-associated and dysbiosis-associated microbial communities across independent cohorts. For example, indole-3- propionic acid, a gut microbiota-derived tryptophan metabolite implicated in intestinal barrier integrity, emerged as one of the strongest health-associated metabolites, whereas imidazole propionate, a dysbiosis-associated metabolite linked to impaired insulin signalling, consistently associated with microbiome dysfunction^49,74,77,79^. Structural similarity analysis further showed that 16 serum and 13 fecal HACK-positive metabolites occupy clinically relevant chemical space shared with approved therapeutics, while experimentally supported microbial producer annotations linked a subset of these metabolites to well-characterized gut bacterial taxa. These annotations carry clear interpretive limits: structural similarity does not imply shared pharmacological activity, and MiMeDB-2.0 associations may represent reported production or transformation potential, whereas monoculture-derived annotations provide experimental evidence of metabolite detection under culture conditions; neither establishes metabolite production or activity *in vivo*. Within those limits, the complementary annotation layers substantially improve biological interpretation and provide a rational framework for prioritizing microbiome-derived metabolites for future mechanistic and translational studies.

Several limitations should be acknowledged. Despite extensive harmonization, some annotations remain unresolved due to incomplete reference spectra or ambiguous structures, and cross-platform comparisons remain influenced by analytical methodology ^85,86^. Our benchmarking also quantifies mapping coverage rather than accuracy; precision benchmarking against a manually curated gold standard remains an important next step. Similarly, microbial producer annotations derived from monoculture experiments cannot fully capture the metabolic complexity of intact microbial communities^26,47^, and continued growth of MiMeDB (CMMC-KB incorporated) and related community-curated resources represents a natural avenue for further strengthening producer attribution within HuMMANet. HuMMANet also harmonizes annotations that already exist and does not assign structures to unannotated spectral features, which remain a substantial fraction of the untargeted metabolome. As additional experimentally validated metabolite resources and microbial annotations become available, these data can be readily incorporated into HuMMANet to further refine producer attribution and biological interpretation.

Beyond its immediate use for cross-study discovery, HuMMANet is well-positioned to support the growing use of foundation models in microbiome science^87–90^. The harmonized microbiome–metabolome pairs assembled here, spanning 54,914 chemically and functionally annotated metabolites linked to pathways, putative microbial producers, physiological distributions, and structural relationships to approved drugs, provide a large-scale standardized corpus for pretraining and fine-tuning metabolite-aware models, directly addressing the annotation heterogeneity that has historically limited such efforts^91,92^. This substantially expands the training and benchmarking corpus available to metabolite prediction and microbe–metabolite association methods such as MelonnPan, mmvec^21^, and MIMOSA/MIMOSA2^18^, which relate microbiome composition to metabolomic profiles and have to date been constrained by the limited scale and heterogeneous annotation of paired training data^93,94^. Looking further ahead, because HuMMANet is organized as explicit relationships linking metabolites to pathways, microbial producers, diseases, and structurally related therapeutics, it can be reformulated as a microbiome–metabolome knowledge graph, supporting link prediction for producer–metabolite and metabolite–disease edges and providing a curated grounding layer for the agentic AI systems now being developed to automate multi-step analytical workflows^95–101^. Future integration with complementary large-scale spectral search infrastructure, such as StructureMASST’s structure-to-sample mapping across public repositories^27^, and microbiomeMASST’s metadata network graph spanning 467 public microbiome metabolomics datasets^28^, could further extend HuMMANet’s producer and physiological-distribution annotations beyond curated study cohorts to the full breadth of public untargeted metabolomics data. More broadly, by providing a standardized metabolite reference onto which future microbiome–metabolome datasets can be directly mapped, HuMMANet establishes a scalable, community-driven resource for reproducible cross-cohort integration, biological interpretation, biomarker discovery, and mechanistic investigation of microbiome-driven host metabolism. Consistent with emerging community-curated microbiome resources such as CMMC-KB^25^, HuMMANet is designed for continuous growth: the accompanying GitHub repository includes a dedicated contribution pathway through which researchers can submit new harmonized studies, updated annotations, or corrections, allowing the resource to expand as the field’s collective knowledge grows.

## Data Integration and Resource Availability

A subset of the microbiome–metabolome datasets included in HuMMANet, including the Lifelines DEEP cohort and the study by Diener *et al.* (2022), was obtained directly from the original study authors under controlled-access agreements. In accordance with the respective data-sharing policies governing these datasets, the associated microbiome and metabolome data are not redistributed through the HuMMANet resource. Researchers wishing to access these datasets should contact the original study authors or follow the data access procedures specified in the corresponding publications or repositories. The HuMMANet computational pipeline, source code, documentation, and associated resource will be made publicly available through an open-access repository upon publication, enabling transparency, reproducibility, and reuse by the research community.

## Acknowledgements

T.S.G. acknowledges the Department of Biotechnology, Ministry of Science and Technology, Government of India for the Ramalingaswami Re-entry Fellowship (BT/HRD/35/02/2006). T.S.G. acknowledges IIIT-Delhi for the Discovery Track Investigator grant. S.V. acknowledges IIIT-Delhi for the Discovery Track Investigator Fellowship.

## Contributions

Conceptualization, T.S.G., S.V., H.M., methodology, formal analysis, validation, T.S.G., S.V., pipeline development, N.A., investigation, T.S.G., S.V., writing-original draft, T.S.G., S.V., writing-review and editing, T.S.G., H.M., supervision, T.S.G., H.M., funding acquisition, T.S.G.

## Corresponding author

Correspondence to Tarini Shankar Ghosh and Himel Mallick.

## Methods

### Study design and scope

This study was designed as a large-scale data integration and resource-building effort to identify, curate, and harmonize publicly available human studies that jointly profile the gut microbiome and metabolome. The primary objective was to construct a comprehensive HuMMANet that enables systematic investigation of microbe–metabolite associations across diverse human cohorts, biological matrices, and disease contexts.

### Identification of microbiome–metabolome studies

Publicly available human microbiome–metabolome studies were identified through a structured search of PubMed, using a combination of controlled vocabulary terms (Medical Subject Headings, MeSH) and free-text keywords. The search strategy was designed to capture studies reporting serum, plasma, fecal, or gut-derived metabolomics data in conjunction with gut microbiome profiling, while excluding non-human and review-only publications. The final Boolean query applied was: (("Humans"[Mesh]) AND ("Metabolomics"[Mesh] OR "Metabolome"[Mesh] OR "Serum Metabolome"[tw] OR "Fecal Metabolome"[tw] OR "Serum Metabolites"[tw] OR "Fecal Metabolites"[tw] OR "Feces"[MAJR]) AND ("Gastrointestinal Microbiome"[Mesh] OR "Microbiota"[Mesh] OR "Microbiome"[tw] OR "Gut Microbiome"[tw] OR "Fecal Microbiota"[tw]) NOT ("Mice"[Mesh]) NOT (Review[Publication Type])) This search retrieved 3,524 publications.

### Screening and eligibility assessment

For all retrieved records, bibliographic metadata including PubMed ID (PMID), digital object identifier (DOI), article title, abstract, year of publication, and journal were extracted. Duplicate records were removed prior to screening. Titles and abstracts were manually reviewed to identify studies reporting integrated, paired, or correlative microbiome and metabolome data derived from human cohorts. Full-text articles were subsequently retrieved and assessed for eligibility according to predefined inclusion and exclusion criteria.

Studies were included if they (i) analyzed human-derived biological specimens, including serum, plasma, feces, saliva, or intestinal biopsies; (ii) reported paired microbiome and metabolome measurements; (iii) provided raw or quantifiable metabolomics data, such as mzXML, mzML, or MGF files, or processed metabolite abundance matrices; and (iv) made data publicly available through established repositories, including NCBI, Metabolomics Workbench, GNPS, ENA, or publisher-linked data portals. Studies were excluded if they were animal-only investigations, review articles or commentaries, or if metabolomics data were unavailable, inaccessible, or insufficiently quantified for harmonization.

### Study inclusion and compendium assembly

Following screening, eligibility assessment, and deduplication, the curated dataset initially comprised 46 distinct cohorts. For harmonization and downstream analyses, these cohorts were further stratified into 59 study units based on analytical modality and data generation strategy, including targeted versus untargeted metabolomics, microbiome sequencing type (16S rRNA gene sequencing or whole-metagenome shotgun sequencing), and biological matrix (serum, fecal, gastric tissue and swab-based metabolomics). This stratification ensured methodological consistency within each study unit while preserving the underlying cohort structure. This stratification was adopted to balance methodological consistency with maximal retention of biological diversity across cohorts.

The complete HuMMANet resource contains multiple biological matrices, whereas the downstream microbiome-association case study was restricted to adult serum and fecal metabolomes. The included studies span a broad range of biological and clinical contexts, including neurological, cardiovascular, metabolic, inflammatory, and gastrointestinal conditions, as well as healthy reference cohorts.

For each included study unit, both study-level and sample-level metadata were systematically extracted and harmonized, including cohort size, disease context, sample type, microbiome profiling strategy, metabolomics platform, sequencing modality, geographic origin, age categories represented, and data accession identifiers. A comprehensive summary of cohort characteristics across all included studies, including sample size, sequencing type, geographic origin, intervention or host exposure context, metabolomics platform and body site, age categories, and the total number of metabolites retained after curation—is provided in **Table S1**.

#### Data extraction and preprocessing

For each eligible study, the following information was curated: study identifier, publication year, DOI, biological matrix analyzed (feces, serum, plasma, saliva, or intestinal tissue), disease condition or cohort classification, reported metabolite names and associated identifiers (PubChem CID, HMDB, KEGG, and ChEBI), analytical platform and acquisition mode (LC–MS, GC–MS, or NMR), and availability of associated microbiome data^41,42,57,58,102^.

Metabolite names lacking biological interpretability or stable chemical identity were excluded — instrumental descriptors (chromatographic modes, ionization states, acquisition platforms), m/z or retention-time encoded names, unnamed placeholder codes (X-prefixed, UNK, ID, peak-based labels), and overly long composite strings — retaining only interpretable, stable identities. Curated data were stored in a unified tabular format with consistent headers and unique study identifiers.

#### Metabolomics data processing

Raw mass spectrometry files, including vendor-specific formats (.wiff, .raw) and open formats (.mzXML), were standardized using MSConvert (ProteoWizard) to generate .mzML files. Untargeted metabolomics datasets were processed using MZmine 3, incorporating peak detection, chromatogram deconvolution, alignment, and feature quantification steps with parameters optimized for centroided data^35,103^. Resulting metabolite abundance tables were exported in .csv format and integrated with curated metadata for downstream harmonization and analysis.

#### Microbiome data processing

Microbiome datasets associated with the included studies were processed using standardized pipelines appropriate to the sequencing strategy employed. For 16S rRNA gene sequencing studies, raw reads were processed using DADA2, including quality filtering, denoising, chimera removal, and inference of amplicon sequence variants (ASVs)^51^. Taxonomic assignment was performed using SPINGO, enabling species-level classification, supported by sequence resolution and reference databases^53^. For whole-metagenome shotgun sequencing studies, microbial taxonomic profiles were generated using MetaPhlAn3, yielding relative abundance estimates at species and genus levels from quality-filtered reads^52^.

### Multi-stage metabolite harmonization pipeline

Reported metabolite names from all studies were harmonized and assigned a unique HuMMANet identifier using a three-stage pipeline. Because metabolite names are frequently reported using inconsistent nomenclature across studies, all annotations were first organized into a unified query metadata table containing the original metabolite name, source study, and HuMMANet identifier. The complete description of this three-stage pipeline is provided in **Text S1** and schematically shown in **Figure 3B**.

#### Stage 1: RefMet-based metabolite standardization

To standardize chemical nomenclature, metabolite names were mapped to the RefMet reference metabolite nomenclature database, which provides curated standardized metabolite names and synonym mappings. RefMet matches were used to retrieve standardized metabolite names together with associated chemical ontology terms, physicochemical descriptors, and database identifiers including PubChem, HMDB, ChEBI, and KEGG.

Metabolites successfully mapped through RefMet were retained as confidence matches, while unresolved metabolites were propagated to subsequent annotation stages.

#### Stage 2: Cross-database identifier reconciliation

Metabolites lacking RefMet mappings were queried against PubChem and HMDB to identify candidate records based on metabolite names and associated synonyms. Matches supported by consistent identifiers across databases were retained as confidence matches, and corresponding chemical annotations were extracted.

Where conflicting annotations were retrieved across databases, HuMMANet retained all candidate records as non-confidence matches and prioritized candidate structures using pairwise structural similarity assessment based on Tanimoto coefficients derived from molecular fingerprints. Metabolites resolved at this stage were annotated with database identifiers and ontology terms, while remaining unresolved metabolites were propagated to the next stage.

#### Stage 3: Name normalization and extended metabolite search

Unresolved metabolites were subjected to name normalization, removing redundant tokens and standardizing query strings. Normalized metabolite names were subsequently queried against the Metabolomics Workbench repository to retrieve additional annotations and database identifiers not captured during earlier stages. Remaining unmatched metabolites were then searched using PubChem offline queries, prioritizing original metabolite names followed by RefMet-standardized names. When matches were identified, HuMMANet retrieved associated chemical ontology classifications, physicochemical descriptors, and database identifiers including PubChem CID, HMDB ID, KEGG ID, and ChEBI ID.

### Integration of biological context

Following harmonization, metabolites were linked to pathway annotations from SMPDB; microbial context from curated MiMeDB-2.0 associations, incorporating CMMC-KB-derived entries, complemented by experimentally derived GNPS/microbeMASST monoculture relationships; and physiological context, including biofluid occurrence and disease associations, from HMDB and related resources. The detailed methodology is schematically described in Figure 3 and Text S1.

#### Benchmarking of metabolite harmonization performance

To benchmark HuMMANet against existing metabolite annotation and mapping frameworks, we used publicly available metabolomics datasets and benchmark datasets previously used for method evaluation. For comparison with MetaboAnalyst, MetaboliteAnnotator, and IDmapping, we used six MetaboLights datasets (MTBLS11733, MTBLS12636, MTBLS13039, MTBLS12997, MTBLS13105, and MTBLS12764) previously used in the evaluation of MetaboliteAnnotator. The corresponding metabolite name lists were used as input to HuMMANet, without additional preprocessing or filtering beyond that applied in the original benchmark. HuMMANet annotation was performed independently for positive- and negative-ionization modes using its deterministic multi-database harmonization framework.

For comparison with metLinkR, we used the five benchmark datasets reported in its original evaluation (Broad 2022, Vicky 2019, DDLPS, LECOC, and COMETS). The standardized metabolite name outputs generated in the metLinkR benchmark were compared with HuMMANet results obtained from the same input metabolite name lists. For all benchmarking datasets, annotation performance was quantified as mapping coverage, defined as the proportion of input metabolite names successfully resolved to at least one standardized metabolite identifier or RefMet entry. No manual curation or post-hoc correction was applied to the HuMMANet outputs, and comparisons were performed using the default or reported configurations of the respective benchmarked methods.

### Cross-study microbiome–metabolome integration using HuMMANet

#### Data preparation and cohort selection

To evaluate the practical utility of HuMMANet in a real-world multi-cohort setting, we conducted a case study integrating serum and fecal metabolomics data across independent adult population cohorts. Analyses were restricted to participants classified as adult, senior, or centenarian to ensure age-appropriate biological comparability across studies. For each biofluid independently, species abundance profiles and rank-scaled metabolome profiles were aligned by matched sample identifiers, retaining only those samples with complete information across species, metabolome, and metadata matrices.

To enable disease-adjusted, study-wise analyses, participant disease status was characterized across clinically defined categories encompassing gastrointestinal, metabolic, cardiovascular, neurological, hepatic, autoimmune, infectious, hematological, renal, respiratory, oral, musculoskeletal, and functional gastrointestinal disorders and encoded as binary indicator variables, with healthy participants serving as the implicit reference category. This encoding ensured that downstream association analyses could statistically account for the confounding influence of disease heterogeneity across cohorts without introducing multicollinearity.

#### Cross-study metabolite retention analysis

To quantify the effect of harmonization on metabolite retention and assess the benefit of HuMMANet harmonization, we evaluated cross-study metabolite retention as a function of progressively stringent detection thresholds defined as the minimum number of independent studies in which a metabolite must be detected for inclusion in downstream analyses.

For each metabolite, detection was determined based on non-zero mean abundance within a given study. Retention curves were constructed by counting the number of metabolites surviving each detection threshold from one to nineteen studies for serum cohorts and one to twenty-two studies for fecal cohorts, separately for unannotated and HuMMANet-harmonized data, and independently for the complete metabolite set and the microbiome-associated subset defined below. Given the wide dynamic range in retained metabolite counts across thresholds, retention values were visualized on a log-transformed scale to facilitate comparison across the full threshold range. For downstream analyses, metabolites detected in at least seven independent studies were retained. This threshold balanced cross-study reproducibility with metabolite retention based on the retention curves shown in **Figure 6A**.

#### Identification of microbiome-associated metabolites

To prioritize metabolites with genuine relevance to gut microbial community structure, as opposed to metabolites that are merely detectable across studies, we applied a permutation-based approach to test each metabolite’s association with overall microbial community composition within each study independently. Specifically, permutational multivariate analysis of variance (PERMANOVA) was conducted using Bray-Curtis dissimilarity between microbial abundance profiles as the response, with each metabolite tested as the sole predictor. Samples with zero total microbial abundance were excluded prior to each test, and missing metabolite values were imputed to zero to retain maximum sample size.

A metabolite was designated as microbiome-associated if it showed a significant association with gut microbial community composition (PERMANOVA, P < 0.05) in at least 50% of the independent studies in which it was detected. This criterion prioritizes metabolites that exhibit reproducible microbiome associations across heterogeneous cohorts while accounting for differences in metabolite detection between studies. The resulting microbiome-associated subset served as the biologically prioritized input for all subsequent HACK score association analyses. *Disease-adjusted species–metabolite association consistency:* To characterize the relationship between individual microbial species and metabolites in a manner robust to disease heterogeneity across cohorts, we developed a disease-adjusted partial Spearman correlation consistency framework. This approach estimates directionally consistent species–metabolite associations across studies while statistically controlling for the confounding influence of disease status.

Within each study, species abundance profiles, metabolome profiles, and disease covariate profiles were combined into a unified matrix for matched samples. Zero-variance columns and rows with missing covariates were removed for numerical stability; rank-scaling infinities were set to missing, with complete-case filtering applied to species and covariates while metabolite missingness was handled via pairwise-complete correlation estimation to preserve sparse metabolomics data.

A second variance filter removed features rendered constant after prior filtering. Partial Spearman correlations across all species–metabolite pairs were then estimated, adjusting for retained disease covariates (or computed as unadjusted correlations where no covariate survived filtering, typically in exclusively healthy or diseased cohorts). Studies with fewer than five samples after all filtering steps were excluded from contribution.

Significance was assessed via t-statistic (degrees of freedom based on contributing observations per correlation), with a directional threshold of p≤0.1 applied per study per pair, and directions recorded and accumulated across studies. The Association-Score was calculated according to the formula given in **Text S3**, and captures predominant direction while penalizing contradictory signals — net directional consistency normalized by per-metabolite detection count, scaled by a penalty proportional to the minority-to-majority directional study ratio. This score ranges continuously from strongly negative to strongly positive, with values near zero reflecting either weak or directionally inconsistent associations across the cohort landscape.

### HACK score correlation and metabolite prioritization

To align metabolite association profiles with the established microbial health axis encoded by HACK scores, the HACK-ranked taxa were matched to the species abundance profiles available in the HuMMANet case-study datasets. Of the HACK-ranked taxa investigated in the original study^48^, 196 overlapped with the species profiles and were therefore included in the analysis. For each metabolite, the Association-Score vector across these HACK taxa was correlated with the corresponding HACK-score vector using Spearman correlation. This yielded a per-metabolite HACK correlation coefficient reflecting the degree to which a metabolite’s cross-study microbial associations are directionally coherent with the health-associated microbial gradient.

To further contextualize each metabolite’s biological relevance, a tri-evidence summary was constructed integrating three complementary dimensions: the proportion of studies in which the metabolite significantly associated with overall microbial community composition, the directional consistency of its association with health-associated species at a predefined significance threshold, and its overall alignment with HACK composite scores across species. Metabolites were ranked by HACK correlation, and representative metabolites from the positive and negative extremes of this ranking were selected for visualization and biological interpretation, reflecting health- and disease-associated microbial metabolic outputs respectively.

#### Sensitivity analysis

To assess the robustness of metabolite-level HACK associations to the potential influence of studies included in the HACK discovery cohort, the HACK score correlation analysis was repeated after excluding these studies. The analysis was performed independently for fecal and serum metabolite datasets using the same analytical framework as the primary analysis. HACK correlation scores from the primary and sensitivity analyses were compared for metabolites shared between the two analyses using Spearman rank correlation. The primary and sensitivity HACK correlation profiles were concordant for both fecal and serum metabolites (**Figure S5**).

#### Pharmacological annotation of HACK-positive metabolites

To investigate the pharmacological relevance of microbiome-associated metabolites, metabolites positively correlated with HACK scores were compared against approved drugs curated from the DrugBank and DrugCentral databases. Canonical molecular structures (SMILES) for metabolites and approved drugs were converted into circular Morgan fingerprints (radius = 2, 2048 bits) using RDKit, and pairwise molecular similarity was quantified using the Tanimoto coefficient. Drug matches identified across DrugBank and DrugCentral were deduplicated using InChIKey prior to downstream summarization, while high-confidence structural analogues (Tanimoto ≥0.75) were used for detailed biological interpretation. DrugBank annotations were further used to assign matched drugs to Anatomical Therapeutic Chemical (ATC) therapeutic categories, enabling characterization of the pharmacological classes represented by structurally similar metabolites. The complete set of drug–metabolite similarities across all thresholds is provided in **Table S16**, while downstream analyses and representative examples were based on the high-confidence subset.

### Functional annotation of microbiome-associated metabolites

Microbiome-associated metabolites identified through the cross-study PERMANOVA framework were functionally characterized using curated pathway, chemical ontology, and microbial producer annotations integrated within the HuMMANet repository. Metabolites that did not satisfy the microbiome-association criterion were retained as the reference group (non-microbiome-associated metabolites) for comparative analyses.

### Pathway enrichment analysis

Metabolic pathway annotations were obtained from the curated Small Molecule Pathway Database (SMPDB) integrated within HuMMANet. Pathway enrichment was evaluated independently for serum and fecal metabolites by comparing microbiome-associated and non-microbiome-associated metabolites. For each pathway, enrichment was assessed using Fisher’s exact test based on the number of microbiome-associated and non-microbiome-associated metabolites assigned to that pathway. Fold enrichment was calculated as:

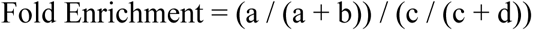

where *a* and *b* represent the numbers of microbiome-associated metabolites within and outside the pathway, respectively, and *c* and *d* represent the corresponding non-microbiome-associated metabolites. P-values were adjusted for multiple testing using the Benjamini–Hochberg false discovery rate (FDR). Pathways with a fold enrichment ≥1.2 and at least four microbiome-associated metabolites were retained for downstream visualization and interpretation.

### Chemical Ontology Enrichment Analysis

Chemical ontology annotations were obtained from the curated ClassyFire classifications integrated within HuMMANet. Enrichment analyses were performed separately for chemical main classes and subclasses using the same statistical framework described for pathway enrichment. Fisher’s exact test was used to compare microbiome-associated and non-microbiome-associated metabolites, followed by Benjamini–Hochberg FDR correction. Chemical classes with fold enrichment ≥1.2 and at least four microbiome-associated metabolites were retained for visualization, while statistical significance was assessed using Fisher’s exact test.

### Species-Level Annotation

Experimentally reported microbial producers were assigned using the integrated MiMeDB-2.0 database together with GNPS monoculture-derived annotations incorporated within HuMMANet. Species annotations from all studies were consolidated into a non-redundant metabolite-to-species mapping by collapsing duplicate species assignments associated with each metabolite. Ambiguous labels, quality-control entries, and non-biological annotations were removed during manual curation. One annotation table was generated for microbiome-associated and non-microbiome-associated metabolites in serum and fecal datasets and was used for biological interpretation.

### Statistical Analysis and Visualization

All statistical analyses were conducted in R (v4.4.1; R Core Team, 2024). Spearman and partial Spearman correlations were calculated using the missing-data handling procedures described above. PERMANOVA analyses were performed using the vegan package with 999 permutations. False discovery rate correction was applied using the Benjamini-Hochberg procedure. All visualizations were generated using ggplot2, with non-overlapping metabolite label placement for volcano plots implemented via ggrepel, and Spearman correlation statistics annotated directly on scatterplots using ggpubr.

