## Supplementary Information (Text S1-S3, Figure S1-S5) for "HuMMANet: A Harmonized Cross-Study Resource for Integrative Analysis of Human Gut Microbiome–Metabolome Associations"

**Complete Methodology of the Harmonization Workflow implemented in HuMMANet**

Metabolomics studies report metabolites using inconsistent, study-specific names, which limits direct comparison and integration across datasets. As study integration increases, the number of metabolites shared across all included datasets declines sharply when identity is inferred from raw metabolite names alone, since the same compound is frequently recorded under different synonyms in different studies (**Figure 3A**). HuMMANet's harmonization framework addresses this by matching each input metabolite name to standardized chemical identifiers; PubChem Compound ID (CID)[1], Chemical Entities of Biological Interest (ChEBI) ID[2], Human Metabolome Database (HMDB) ID[3], and Kyoto Encyclopedia of Genes and Genomes (KEGG) ID[4]; through a three-stage matching procedure of increasing scope, followed by automated biological and pharmacological annotation (Figure 3B). Each metabolite retains a unique internal identifier and its original input name throughout the workflow, enabling full traceability of results. Collectively, the pipeline reconciled 33,469 study-specific annotations into 14,550 unique metabolites (13,063 Annotated by this harmonization framework). (**Figure 3B**).

***Stage 1: Standardization against RefMet:*** Input metabolite names from all integrated studies are first queried against an in-house curated synonym library for direct identification (Figure 3B). This library comprises 54,914 unique metabolite names and their associated synonyms, aggregated from the 59 integrated studies used in this analysis and fixed as a static reference resource; each newly incoming study's metabolite names are searched against this frozen set of names and synonyms. Expansion of the library is planned for future versions of the workflow. Names not resolved by this initial lookup, including slash-delimited composite entries that are first parsed into individual candidate names, are queried against the RefMet nomenclature database[5], which returns a standardized name, known synonyms, and cross-referenced identifiers where available. Returned PubChem CIDs are validated against a locally maintained offline PubChem database and supplemented using the PubChem PUG-View application programming interface (API) to recover any missing HMDB, KEGG, or ChEBI identifiers associated with that CID. Records still lacking identifier information are further supplemented from a curated local HMDB reference table, including a provisional chemical classification (superclass, class, and subclass) drawn from a local PubChem classification table. A metabolite is considered mapped at this stage if it is associated with at least one PubChem CID, ChEBI ID, HMDB ID, or KEGG ID; all remaining unmapped metabolites proceed to Stage 2 (**Figure 3B**).

***Stage 2: Cross-Database Reconciliation:*** Metabolites unresolved by Stage 1 are queried independently against local HMDB and PubChem name-matching indices and against the PubChem PUG-View API. For each query, HuMMANet retrieves (i) HMDB identifiers and any PubChem CID recorded within HMDB, and (ii) PubChem CIDs matched by name and any HMDB identifier cross-referenced to that CID. Queries for which both sources return concordant identifiers, and for which each source returns a single unambiguous candidate identifier, are retained as confidence matches; queries for which the two sources disagree, for which only one source returns a result, or for which either source returns more than one candidate identifier for the same query, are retained as non-confidence matches, with one row generated per candidate identifier combination (Figure 3B). Queries still unmatched after this initial pass are re-queried a second time with trailing wildcard characters stripped from the input name, to recover matches lost to minor formatting artifacts. No candidate is discarded or resolved by default at this stage.

For each candidate, the corresponding structure is retrieved (as a Simplified Molecular Input Line Entry System, SMILES, string) from HMDB or PubChem and compared by Tanimoto similarity, computed from 2048-bit Morgan fingerprints (radius = 2, RDKit), against a reference structure selected within its identifier group, prioritizing HMDB-derived structures where available and falling back to any candidate with a valid structure otherwise. Each candidate structure is also annotated with chemical ontology terms (superclass, class, subclass) using ClassyFire[6], queried once per unique International Chemical Identifier Key (InChIKey) against the public ClassyFire web service via the classyfireR R package[7], accessed from the Python pipeline through an rpy2 bridge; queries that fail or return no classification are left unannotated rather than causing the pipeline to halt. This step provides a quantitative basis for downstream evaluation of competing candidate matches. Metabolites without a resolved identifier proceed to Stage 3.

***Stage 3: Extended Search and Recovery*****:** Remaining unmatched names are first queried by name against the Metabolomics Workbench reference database[8]; where an existing record is found, its associated KEGG, PubChem, and HMDB identifiers are retrieved directly (Figure 3B, Stage 3A).

Names without an existing Metabolomics Workbench record undergo additional normalization, including standardization of Unicode dashes and punctuation, whitespace collapsing, generation of alternate candidate strings for conjunction-separated names (e.g., "X or Y"), and removal of lipid-shorthand parenthetical suffixes, to maximize the likelihood of exact-string matching (Figure 3B, Stage 3B). Normalized names are then re-queried against Metabolomics Workbench, including its table of previously unmapped metabolite names, and, where enabled, against a reverse synonym-to-CID index derived from the local PubChem database. Names remaining unresolved are passed through the RefMet bridge a second time as a final matching step. Any PubChem CID obtained through these routes is enriched using local property tables (SMILES, InChIKey, standardized name, International Union of Pure and Applied Chemistry [IUPAC] name, synonyms), a live PubChem properties API call, PUG-View identifier backfill, and ClassyFire annotation. Metabolites with at least one identifier (PubChem CID, HMDB ID, KEGG ID, or ChEBI ID) after this stage are considered mapped; all others are reported as unresolved.

***Biological Annotation*: ** Following each stage, all newly mapped metabolites are passed to a shared annotation module that appends metabolic pathway associations via SMPDB[9], curated and experimentally derived microbial producer associations via MiMeDB 2.0[10] and GNPS/microbeMASST[11,12]; physiological associations, including biofluid distributions and disease associations, via HMDB[3]; and pharmacological annotations via DrugBank[13] and DrugCentral[14] to that stage's output.

***Pharmacological Annotation*: ** For each mapped metabolite with a resolvable structure, the canonical SMILES string is processed using RDKit[15] to generate a chirality-aware Morgan count fingerprint (radius = 2) — distinct from the achiral, bit-vector Morgan fingerprint used for candidate-structure comparison in Stage 2. This fingerprint is compared, via bulk Tanimoto similarity[16], against two independently prepared drug reference libraries. DrugBank structures were obtained from its structure data file (SDF) and cross-annotated with metadata parsed from the DrugBank XML export, including Anatomical Therapeutic Chemical (ATC) codes[17], ATC classification text, clinical indication, and mechanism of action; ATC classification text was further mapped to a coarse therapeutic-area label (e.g., cardiovascular, neurological, oncological, respiratory, dermatological) by keyword matching against standard top-level ATC categories. DrugCentral structures were obtained from its structure and identifier export and converted directly to molecular objects from the provided SMILES[18]; entries whose SMILES could not be parsed into a valid molecular structure were excluded from the reference library. For each metabolite, the five most structurally similar compounds from each reference library are retained, together with the Tanimoto similarity score, compound identifier and name, and, for DrugBank matches, ATC code, ATC class, indication, mechanism of action, and assigned therapeutic area; no minimum similarity threshold is applied, so the five best-available matches are reported even when overall structural similarity is low.

***Package Design and Reproducibility*****:** The workflow is implemented as independently executable stage modules that share a common set of local reference resources: an offline PubChem database (rebuilt from public PubChem source files rather than distributed as a static file, to minimize package size while preserving reproducibility), a curated local HMDB reference table, the Metabolomics Workbench database export, and the DrugBank and DrugCentral structure libraries used for pharmacological annotation. Each stage can be executed independently or as part of a chained pipeline. Mapped, unmapped, and non-confidence match tables, along with corresponding output workbooks, are written to disk after every stage, allowing intermediate inspection, resumption from any point, and re-analysis of unresolved metabolites in subsequent stages. Reference paths are resolved relative to the installed package location to support consistent execution across computing environments. Two annotation steps (PubChem PUG-View identifier backfill and ClassyFire chemical ontology lookup) query live external web services rather than local snapshots; because the underlying source databases are updated independently of this package, results from these steps may vary slightly between runs performed at different times, even when local reference files and code are unchanged.

***Data Outputs:*** All outputs are provided in tabular format for downstream analysis, including mapped metabolites, confidence matches, non-confidence matches, unresolved metabolites, and integrated annotation workbooks containing pathway, microbial producer, physiological, disease, and pharmacological information. Pharmacological outputs include structurally similar approved drugs, molecular similarity scores, ATC classifications, clinical indications, mechanisms of action, and therapeutic-area assignments. All tables retain both the original, study-specific metabolite name and the corresponding matched identifiers and annotations, supporting full traceability of the harmonization process.

#### Text S2

##### Integration of the MicrobeMASST Monoculture Collection into HuMMANet

To expand the microbial coverage of the HuMMANet knowledgebase, we generated a comprehensive, harmonized reference of microbially produced metabolites derived from the MicrobeMASST monoculture collection[11], a public repository of mass spectrometry data acquired from single-organism ("monoculture") microbial cultures. For each metabolite detected, we identified its likely microbial source and its co-occurrence with human biofluids. Metabolite identification was performed using GNPS-based molecular networking (defined below)[12,19], and the resulting microbe–metabolite associations were harmonized into standardized identifiers using the HuMMANet workflow (**Text S1**) before integration into the resource. The overall process comprised five stages: (i) automated retrieval and preparation of monoculture metadata and molecular networking submission; (ii) processing of molecular networking results; (iii) integration and consolidation of outputs across species; (iv) blank and media subtraction for quality control; and (v) aggregation and biofluid annotation of the resulting metabolite profiles. Each stage is described in detail below.

***Background: GNPS and Molecular Networking*****:** The Global Natural Products Social Molecular Networking platform (GNPS) is a public, web-based infrastructure for the analysis, sharing, and spectral library matching of tandem mass spectrometry (MS/MS) data[19]. Molecular networking is an analytical approach implemented within GNPS that groups MS/MS spectra according to their spectral similarity, on the principle that structurally related metabolites tend to produce similar fragmentation patterns. Each group of related spectra forms a molecular "cluster," and each cluster is compared against reference spectral libraries to assign a putative chemical identity. Applying molecular networking to monoculture mass spectrometry data therefore allows the metabolites produced by a given microbial species to be identified from raw spectral data in a systematic and reproducible manner. The MassIVE data repository (Mass Spectrometry Interactive Virtual Environment) is the data-storage system underlying GNPS submissions, and each deposited dataset is assigned a unique MassIVE identifier[19].

***Retrieval of Monoculture Metadata from MicrobeMASST*****:** Sample- and organism-level metadata for all microbial monoculture datasets were obtained from the MicrobeMASST resource[11], which catalogues publicly deposited monoculture mass spectrometry datasets together with their associated taxonomic and experimental annotations. This metadata was consolidated into a master metadata table (microbe_masst_table_metadata) containing, for each sample, its source organism, taxonomic identifiers, MassIVE dataset identifier, and associated raw data filenames. This table served as the single input from which all subsequent organism-specific processing and GNPS submissions were generated programmatically, ensuring that every downstream molecular networking job was linked back to a defined and traceable set of monoculture samples.

***Automated GNPS Submission Framework*****:** To enable molecular networking analysis of monoculture metabolomics data at scale, we developed an automated GNPS submission framework implemented in Python. The framework performs structured metadata preprocessing and automates interaction with the GNPS web platform, enabling high-throughput submission of large numbers of MassIVE datasets while ensuring consistent workflow configuration across all analyses. This automation was necessary because the MicrobeMASST monoculture collection spans thousands of individual datasets, and manual submission and configuration of each dataset through the GNPS web interface would not be feasible at this scale.
The pipeline is organized as a modular architecture comprising four coordinated scripts (FinalDestination.py, ip3.py, ip10.py, and ip11.py), each responsible for a distinct stage of metadata preparation, workflow configuration, and job submission.

***Metadata preprocessing and organism-level structuring*****:** The master metadata table is parsed using the Pandas library[20,21]. Organism identities are extracted from the attribute_taxaname_file field, and entries are grouped accordingly to generate organism-specific metadata files in tab-separated (.tsv) format. For each sample entry, an Attribute_group field is constructed using conditional logic based on the attribute_taxa_NCBI column, explicitly preserving quality control ("QC") and blank samples as distinct categories while assigning all remaining entries to their corresponding organism labels — this preservation of blank and QC labels at the metadata stage is what subsequently allows these samples to be identified and excluded during blank subtraction (see below). Each organism-specific metadata file is formatted for compatibility with GNPS molecular networking submission requirements.

***Automated GNPS workflow execution*:** Following metadata generation, GNPS workflow execution is automated using Selenium-based browser control, a software approach that programmatically drives a standard web browser to interact with a website exactly as a human user would, but without manual input. Execution is initiated through FinalDestination.py, which sequentially orchestrates three operational modules:

- ip3.py establishes an authenticated GNPS session by navigating the ProteoSAFe interface[19,22], submitting user credentials, and positioning the session within the Molecular Networking workflow[12], with explicit synchronization steps to ensure reliable interaction with dynamically loaded page components.
- ip10.py automates interaction with the GNPS workflow configuration interface: organism-specific metadata files are programmatically assigned to the required spectrum file input, and molecular networking parameters — precursor and fragment ion mass tolerances, cosine similarity thresholds, minimum fragment ion counts, and minimum matched-peak requirements for spectral library searches — are configured identically across all submissions to ensure comparable results between species. Structured exception handling maintains stability against variability in the GNPS front-end interface.
- ip11.py governs batch-level dataset processing and submission management: MassIVE dataset identifiers are extracted from the metadata inputs, and datasets are processed sequentially to prevent a failure in one submission from affecting others. Session-management routines automatically re-establish authentication if interrupted, and browser sessions are terminated in a controlled manner after each successful submission to prevent resource leakage.

**Scalability, robustness, and reproducibility.** All jobs are assigned structured, metadata-derived identifiers and executed in batch mode, enabling molecular networking to be run across thousands of microbial monoculture datasets with minimal manual intervention. Logging and diagnostic outputs are retained for every job to support reproducibility and downstream verification. All automation scripts, metadata templates, and processed outputs are publicly available through the accompanying GitHub repository, enabling transparent reuse, inspection, and independent reproduction of the molecular networking analyses.

***Retrieval and Processing of Molecular Networking Results*****:** Species-specific GNPS molecular networking results: The output of the process described above were collected and harmonized into a single, study-agnostic reference dataset describing microbe–metabolite associations. For each microbial species, GNPS output tables corresponding to molecular cluster annotations, cluster membership ("bucket") tables, spectral annotations, and experimental metadata were retrieved directly from the compressed GNPS result archives without requiring manual file extraction. Only tables that were successfully parsed and non-empty were retained for downstream processing. Species-level metadata tables submitted during GNPS analysis were matched back to molecular features using filename-based identifiers, ensuring a consistent link between each detected metabolite feature and the experimental sample it originated from. To maintain completeness across closely related datasets, metadata were propagated across replicate or derivative entries of the same species where appropriate, allowing uniform annotation of molecular features originating from the same biological source.

***Integration and Consolidation Across Species*****:** GNPS-derived annotation tables, cluster membership tables, and experimental metadata were integrated for each species using a standardized processing function. Molecular features were represented at the level of GNPS cluster indices, which correspond to consensus molecular features derived from grouping spectrally similar MS/MS scans. Processing across the large number of species in the collection was performed in batches with checkpointing, allowing recovery from partial failures without needing to reprocess previously completed species and ensuring reproducibility of the full pipeline. Duplicate species entries arising from repeated processing runs were explicitly identified and resolved, retaining a single representative dataset per species. The resulting species-resolved molecular profiles contain metabolite feature intensities aggregated across all samples and files belonging to that species, enabling downstream comparative analysis across microbial taxa.

***Blank and Media Subtraction*:** To remove background signals originating from culture media, blank injections, or other technical controls rather than the microorganism itself, a conservative blank-subtraction procedure was applied to every species-specific molecular profile. For each molecular feature (GNPS cluster), the maximum signal intensity observed across all blank samples was compared to the maximum signal intensity observed across all biological (monoculture) samples. A feature was retained only if its biological-sample intensity exceeded its blank-sample intensity by at least three-fold; features not meeting this threshold were removed as likely media- or background-derived signals. In addition, any molecular features linked, via their sample metadata, specifically to blank, media-only, or quality-control filenames were excluded outright, independent of the intensity-ratio criterion. This two-part filtering strategy ensured that retained molecular features represented metabolites reproducibly and specifically detected in microbial monocultures rather than experimental artifacts introduced by the growth medium or sample handling. Post-filtering validation confirmed complete removal of blank-, media-, and QC-associated features across all species in the collection.

***Aggregation of Metabolite Profiles Across Taxa*:** Following blank subtraction and quality control, metabolite profiles from all microbial species were combined into a single unified dataset. For each metabolite–species pair, metabolite intensities were summarized using total, mean, and maximum intensity across samples, together with the frequency with which the metabolite was detected across samples and files. Metabolites were grouped using stable chemical identifiers where available, including InChIKey, compound name, and cross-referenced database identifiers, to ensure that molecular clusters representing the same underlying chemical entity, even if detected in multiple separate GNPS jobs, were correctly consolidated rather than counted as distinct metabolites. This produced a compact, biologically interpretable table of metabolite production across microbial taxa. A binary presence–absence matrix was additionally constructed, recording whether each metabolite was detected in each microbial species, to support downstream network analyses and integration with human cohort data.

***Annotation of Metabolite Biofluid Origin:*** To place the identified microbial metabolites in the context of human physiology, each detected metabolite was annotated according to its reported occurrence in human biofluids, using a curated subset of the Human Metabolome Database

(HMDB)[3]. HMDB records were collapsed by InChIKey to generate a single, unified mapping between metabolites and their associated human biofluids. Metabolites were classified as "bacterial-only" (not reported in any human biofluid) if no overlap with major human biofluids was found, or as "bacterial and human-associated" if detected in one or more of serum, urine, feces, cerebrospinal fluid, saliva, or sweat. Metabolites lacking a resolvable chemical identifier were conservatively labeled as having unknown biofluid origin rather than being assigned to either category. Where available, the specific biofluids associated with each metabolite were retained as structured annotations, enabling fine-grained interpretation of which microbial metabolites may be relevant to human physiology.

***Final Resource Assembly and Availability*: **All processed outputs from the microbial monoculture molecular networking analysis, including metabolite annotations, microbial source associations, biofluid origin labels, chemical classifications, and quality-controlled abundance summaries, were harmonized into standardized identifiers via the HuMMANet workflow (**Text S1**) and integrated into the HuMMANet resource. These data are provided as study-agnostic, analysis-ready tables, enabling direct reuse and independent validation. The complete set of processed monoculture-derived microbe–metabolite associations, together with all harmonized annotations and metadata, is publicly available through the accompanying GitHub repository, supporting transparent and reproducible downstream analyses.

#### Text S3

##### Identification of HACK-associated microbiome-derived metabolites

To determine whether microbiome-associated metabolites preferentially associate with microbial taxa linked to host health, we integrated the microbiome association score (MAS), metabolite–taxon association score (AS), and the Health-Associated Core Keystone (HACK) framework (**Figure S3**)[23].

Microbiome-associated metabolites (MAS ≥ 0.5) detected in at least four studies were analyzed separately for serum and fecal datasets. Only metabolites detected in more than seven studies were retained for the final HACK association analysis reported in the Results. For each metabolite, its association with every HACK-ranked microbial taxon was quantified in two steps.

First, metabolite–microbiome associations were evaluated independently within each study. Bray–Curtis distance matrices were calculated from species abundance profiles, and PERMANOVA was performed to test the association between microbiome composition and each metabolite[24]. Studies with $P < 0.05$ were considered significant. For each metabolite, the microbiome association score (MAS) was calculated as the proportion of studies showing a significant association among all studies in which that metabolite was detected. Metabolites with MAS ≥ 0.5 were retained for subsequent analyses.

Next, partial Spearman correlations were calculated within each study between the abundance of each retained metabolite and each HACK taxon while adjusting for study-specific confounding variables (disease variables). For every metabolite–taxon pair, these correlations were summarized across studies using the Association Score (AS):

$$AS_{ij} = \frac{SP_{ij} - SN_{ij}}{N_{\text{detected},j}} \left(1 - \frac{\min(SP_{ij}, SN_{ij}) + 1}{\max(SP_{ij}, SN_{ij}) + 1}\right)$$

where $SP_{ij}$ and $SN_{ij}$ denote the number of studies showing significant positive and negative partial Spearman correlations ($P \leq 0.1$), respectively, between metabolite $j$ and taxon $i$, and $N_{\text{detected},j}$ is the total number of studies in which metabolite $j$ was detected. The first term quantifies the overall direction of association across studies, whereas the second term penalizes metabolite–taxon pairs with similar numbers of positive and negative associations. The resulting AS positive values indicate predominantly positive associations across studies and negative values indicates predominantly negative associations. This generated an association profile for each metabolite across HACK taxa.

Finally, each metabolite's AS profile was correlated with the HACK scores of all HACK taxa (predefined from original HACK paper) using Spearman correlation[23]. Metabolites showing significant positive correlations (P value $\leq 0.05$) were classified as HACK-positive microbiome-associated metabolites, whereas those showing significant negative correlations (P value $\leq 0.05$) were classified as HACK-negative microbiome-associated metabolites.

#### Figure S1

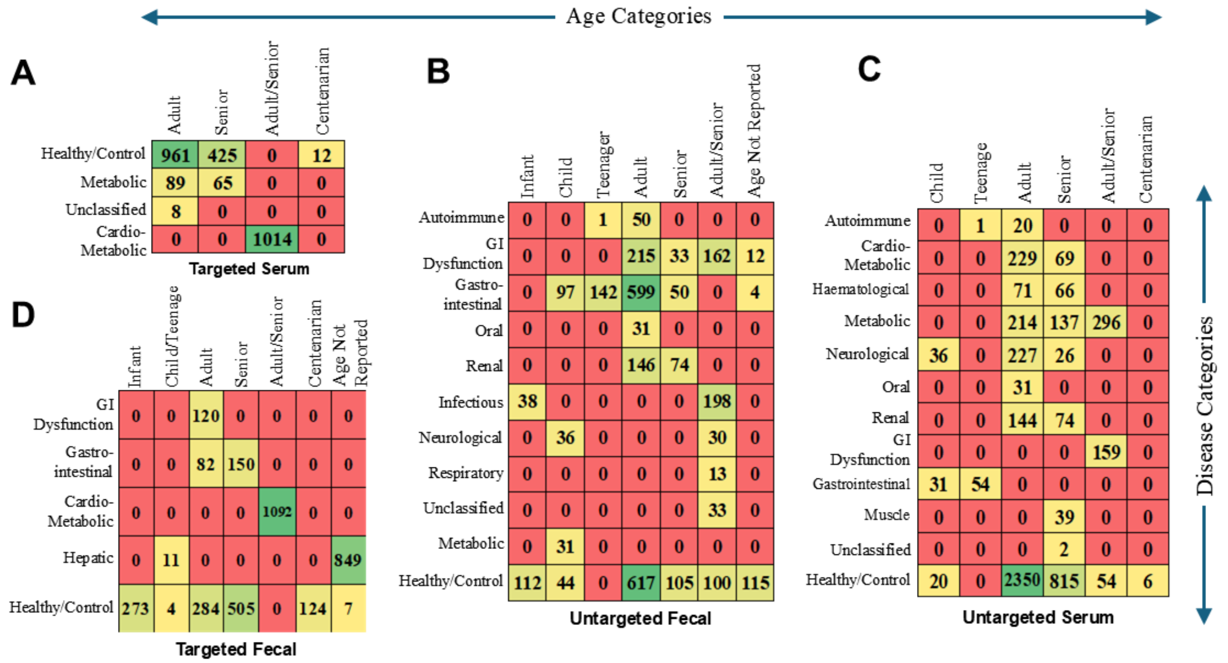


**Figure S1: Distribution of HuMMANet samples across disease and age categories, stratified by metabolomics platform and sample type.** Heatmaps show the number of samples in the HuMMANet dataset across disease category (rows) and age category (columns) combinations, separately for four platform–matrix strata: **(A)** targeted serum, **(B)** untargeted fecal, **(C)** untargeted serum, and **(D)** targeted fecal metabolomics. Age categories (top) range from infant through centenarian, including composite (Adult/Senior, Child/Teenage) and unreported categories where applicable; disease categories (right) include Healthy/Control alongside clinically defined conditions such as Metabolic, Cardio-Metabolic, Gastrointestinal, Autoimmune, Neurological, Renal, Hepatic, Haematological, Respiratory, Oral, and Musculoskeletal disorders, among others. Each cell reports the raw sample count for that disease–age combination, with colour intensity scaled by count within each panel (red, low; yellow, intermediate; green, high). Category composition and count ranges vary across the four panels, reflecting differences in cohort representation between targeted and untargeted platforms and between serum and fecal biofluids.

**Figure S2**

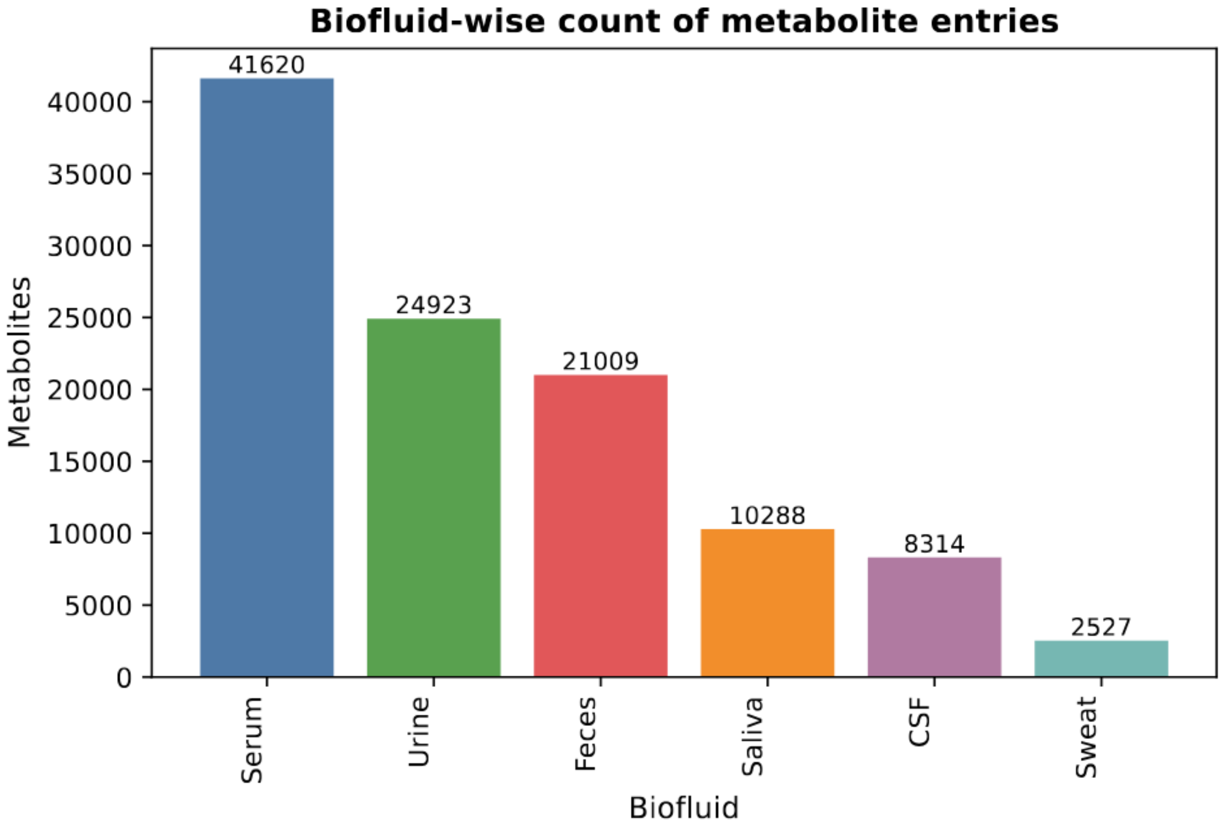


**Figure S2: Distribution of GNPS derived annotated metabolites across human biofluids.** Bar plot showing the number of unique metabolites annotated in major human biofluids based on Human Metabolome Database (HMDB) provenance. Metabolites detected in serum, urine, feces, saliva, cerebrospinal fluid (CSF), and sweat are shown, highlighting the broad physiological coverage of microbially derived and human-associated metabolites integrated into HuMMANet. Counts reflect unique metabolites with resolvable chemical identifiers and biofluid annotations.

### Figure S3

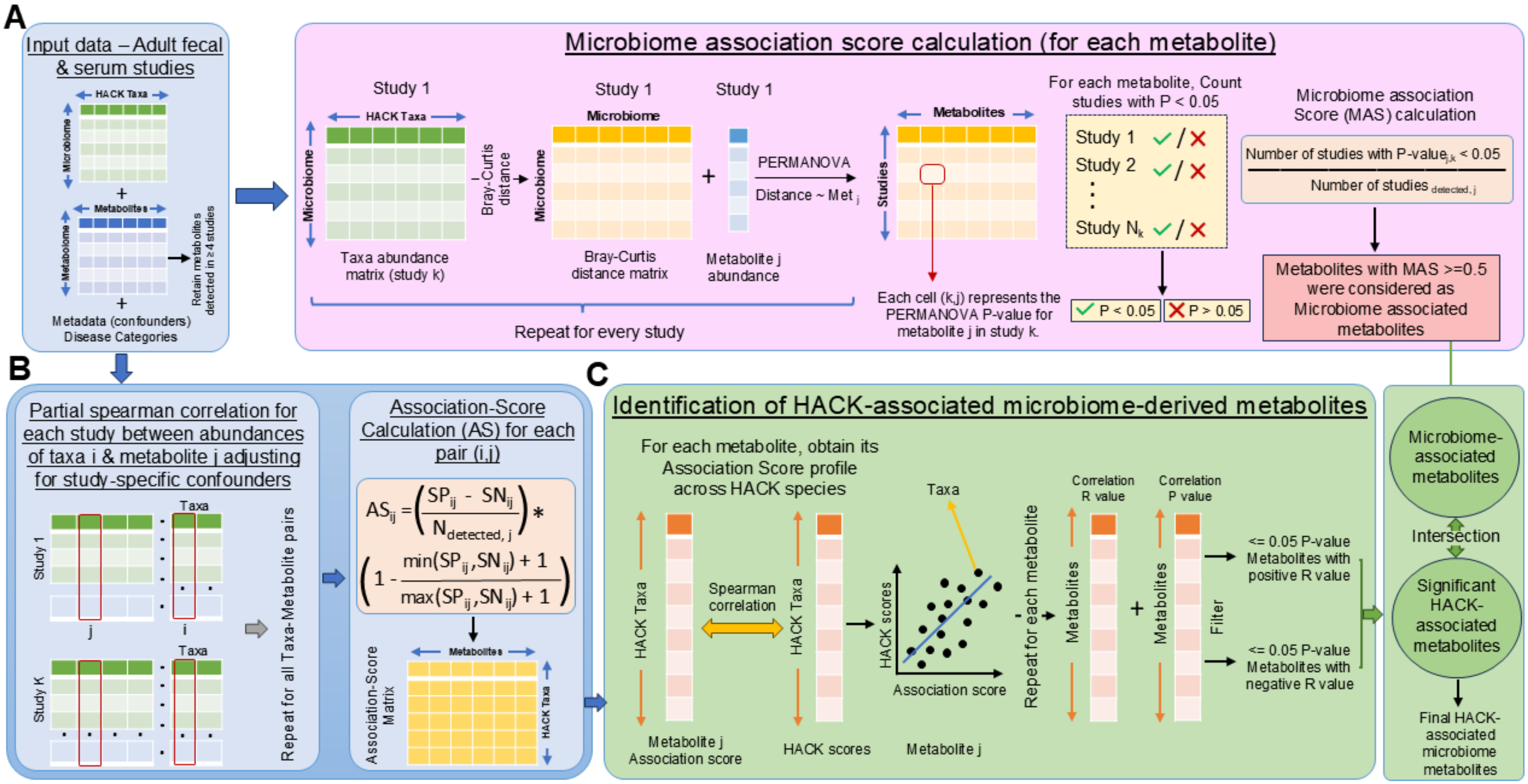


$SN_{ij}$ = number of studies where partial spearman correlation is -ve with at least p<=0.1
$SP_{ij}$ = number of studies where partial spearman correlation is +ve with at least p<=0.1
$N_{detected,j}$ = Total studies where metabolite j was detected

HACK-positive microbiome metabolites = MAS ≥ 0.5, positive HACK correlation (P ≤ 0.05)
HACK-negative microbiome metabolites = MAS ≥ 0.5, negative HACK correlation (P ≤ 0.05)

**Figure S3: Workflow for identifying microbiome-associated metabolites and HACK-associated metabolite signatures (A) Identification of microbiome-associated metabolites.** Adult paired microbiome–metabolome datasets were harmonized using HuMMANet. For each study, Bray–Curtis distance matrices were calculated from microbial taxonomic profiles, restricted to the HACK-taxa[23]. Associations between microbial community composition and individual metabolite abundances were assessed using PERMANOVA, adjusting for study-specific covariates. For each metabolite, a microbiome association score (MAS) was defined as the proportion of studies in which the metabolite showed a significant association with microbial community composition ($P < 0.05$); metabolites with MAS $\geq 0.5$ were classified as microbiome-associated. **(B) Computation of Association-Scores between metabolites and the HACK microbial taxa.** Partial Spearman correlations were computed between the levels of each metabolite with the HACK-taxa within each study, adjusting for study-specific covariates. For each taxon-metabolite pair, an Association-Score (AS) was calculated by integrating the consistency and direction of these associations across studies, generating an Association-Score profile for each of the metabolites. **(C) Identification of HACK-associated microbiome-derived metabolites.** For each microbiome-associated metabolite (from A), Spearman correlations were calculated between the Association-Score profile of the metabolite with the different HACK-taxa and the taxon-specific HACK-scores across samples. Metabolites with Association-Score profiles showing significant positive or negative correlations with taxon-specific HACK scores ($P \leq 0.05$) were intersected with the microbiome-associated metabolite set (from A) to define the final set of HACK-associated microbiome-derived metabolites.

**Figure S4**

**A HuMMANet Improves Cross-study Metabolite Retention In Serum Adult Cohort**

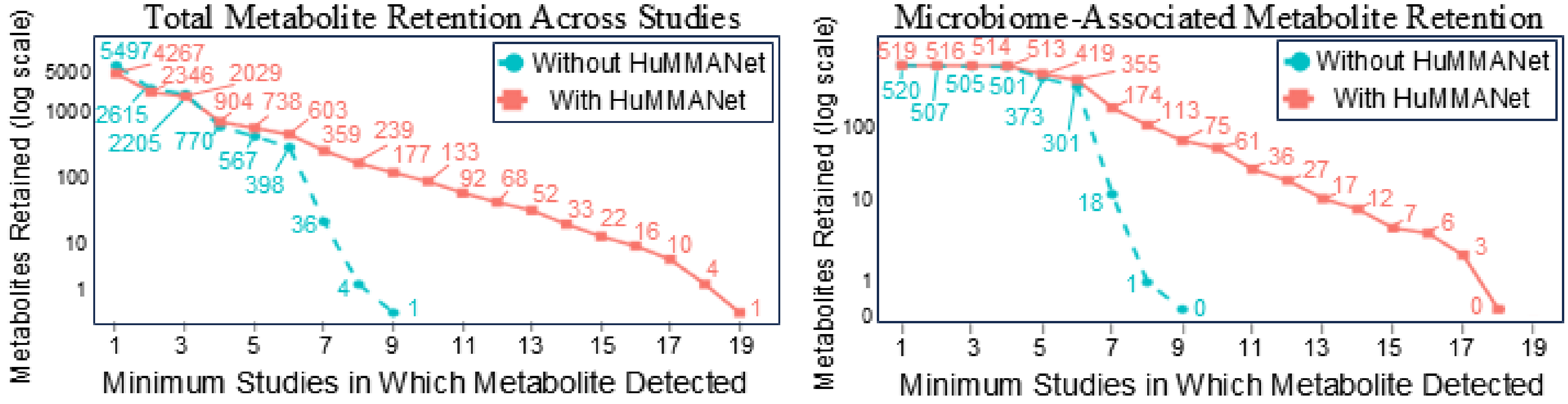


**B HuMMANet Improves Cross-study Metabolite Retention In Fecal Adult Cohort**

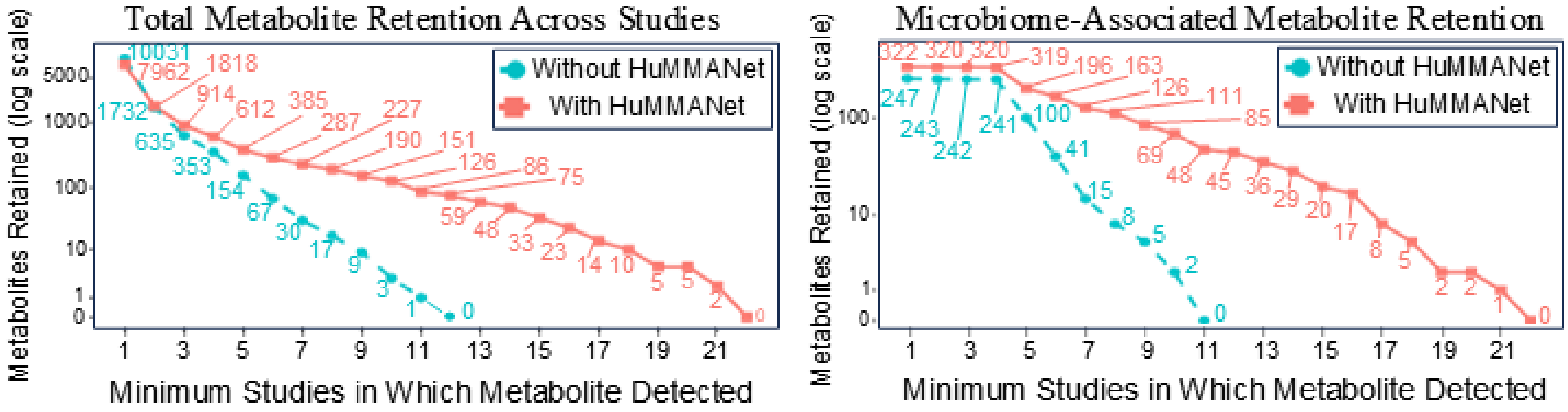

**Figure S4: HuMMANet Enables Cross-Study Microbiome-Metabolite Integration.**

Cross-study metabolite retention curves, shown as a function of the minimum number of studies in which a metabolite must be detected for inclusion (x-axis), with and without HuMMANet harmonization (teal: without; pink: with). Y-axis: number of metabolites retained (log1p scale). Left panels show retention across all detected metabolites; right panels show retention restricted to microbiome-associated metabolites (PERMANOVA, $P < 0.05$, in ≥1 study). **(A)** Serum adult cohorts. **(B)** Fecal adult cohorts. HuMMANet substantially increased retained metabolite counts at every threshold in both biofluids, particularly among microbiome-associated compounds.

**Figure S5**

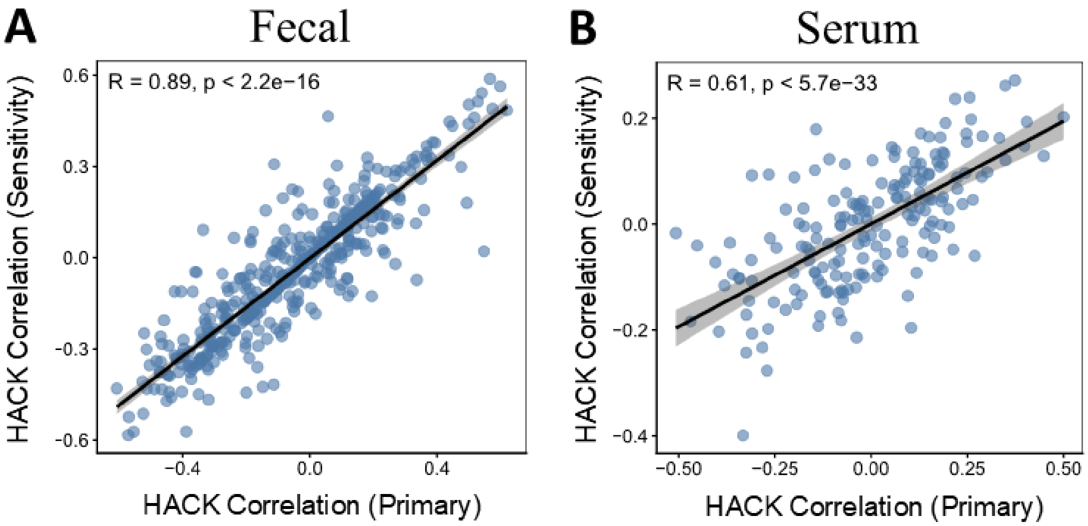


**Figure S5: Sensitivity Analysis of Metabolite-Level HACK Correlation Scores. (A)** Comparison of metabolite-level HACK correlation scores obtained from the primary analysis and from a sensitivity analysis performed after removing studies included in the HACK discovery cohort, for fecal metabolites. Each point represents a metabolite shared between the two analyses, with the fitted regression line and corresponding Spearman rank correlation coefficient and P-value shown. **(B)** Corresponding comparison for serum metabolites. In both panels, each point represents a metabolite shared between the two analyses, and the fitted regression line is shown.
